# Genomic prediction of disease resistance provides a path to marker assisted restoration in a wetland foundation tree species

**DOI:** 10.1101/2025.09.14.675737

**Authors:** Karina Guo, Collin Ahrens, Stephanie Chen, Karanjeet Sandhu, Maurizio Rossetto, Ashley Jones, Chloe Tan, Justin Borevitz, Richard Edwards, Jason Bragg

**Affiliations:** Research Centre for Ecosystem Resilience, Botanic Gardens of Sydney, NSW, Australia; School of Life and Environmental Sciences, The University of Sydney, NSW, Australia; Evolution & Ecology Research Centre, School of Biotechnology and Biomolecular Sciences, UNSW Sydney, Sydney, NSW, Australia; Cesar Australia, Brunswick, VIC, Australia; Research School of Biology, The Australian National University, Canberra, ACT, Australia

**Keywords:** myrtle rust, *Melaleuca quinquenervia*, GWAS, genomic prediction, restoration

## Abstract

Tree species worldwide are under threat from non-native pathogens that impact forests and the ecosystem services they provide. Myrtle rust, caused by *Austropuccinia psidii*, is one example, first detected in Australia in 2010. This fungal pathogen infects immature tissue from a wide range of Myrtaceae hosts, including the wetland foundation species *Melaleuca quinquenervia*. Durable restoration action for this species would preferentially incorporate disease resistant individuals. Our aim for this study was to provide a molecular assay to identify and select resistant individuals and seed lots for restoration. We conducted artificial inoculation of a panel of seedlings and measured their immune responses to myrtle rust. We then performed whole genome sequencing (3.2M common SNPs) and conducted a Genome Wide Association Study (*N* = 492), which revealed clusters of significantly associated SNPs in three chromosomal regions, including clusters of putative R genes. Associated SNPs were filtered to a panel of 1,049 for a highly accurate genomic prediction model (*R* = 0.83). This provides a relatively inexpensive approach to identifying resistant individuals or seed lots for restoration and a template for managing myrtle rust impacts while maintaining population genetic diversity.

## Introduction

Exotic plant diseases are a global threat to conservation, ecosystems, and economic security (Ristaino et al., 2021; Jaureguiberry et al., 2022). The damage caused by pathogen incursions can lead to population contractions (e.g., Cunningham et al., 2021; Luedtke et al., 2023), threatens species (Fensham & Radford-Smith, 2021), and impacts the function and composition of ecosystems (Anagnostakis, 1987; Smith et al., 2006; Stevenson et al., 2023). In agriculture, invasive pathogens can cause large-scale crop loss (Savary et al., 2019). These impacts are expected to accelerate with predicted changes in climate (Singh et al., 2023).

*Austropuccinia psidii* (G.Winter) Beenken, which causes the disease myrtle rust, is a devastating global exotic plant fungus known to infect species across a broad range of genera in the Myrtaceae family (Soewarto et al., 2018). It was first detected in Australia in 2010 and poses a major threat to Australian forest ecosystems that are dominated by Myrtaceae (Carnegie & Cooper, 2011). The disease targets soft tissue, including immature leaves, buds, fruits, and branches (Carnegie et al., 2016; Fernandez-Winzer et al., 2019; Makinson, 2018; Sandhu & Park, 2013). These impacts vary substantially among the Australian Myrtaceae, from species that are unaffected by myrtle rust (Sandhu & Park, 2013), to others that are highly susceptible and face severe population contractions due to the disease (Makinson, 2018).

Consequently, several species have been listed as Critically Endangered in Australia (Threatened Species Scientific Committee 2020a, 2020b), and the impacts of myrtle rust are recognised as part of a ‘threatening process’ under relevant legislation in several Australian jurisdictions (for instance the *Environmental Protection and Biodiversity Conservation Act 1999*).

The protection of species and ecosystems that are impacted by myrtle rust is challenging. For several highly imperilled species, collections have been gathered and are maintained ex situ, protected by fungicide (Pathan et al., 2020; Hardstaff et al., 2025). Ideally, these populations would be reintroduced to natural sites in self-sustaining populations, but this would require breeding programs to increase resistance to the pathogen (Makinson, 2018). Fungicides can rarely be applied in wild reserves, due to potential risks of toxicity (Shen et al., 2024) and impacts on other fungi (Zimmer et al., 2023). In some species, there is variation in resistance to myrtle rust among individuals. For instance, in *Eucalyptus* this variation is observed to be heritable (Miranda et al., 2013). As a result, breeding programs for genetically resistant individuals have been successfully implemented in forestry. A similar approach could be used in natural populations with the introduction or promotion of resistant genotypes to help mitigate the disease’s impact on ecosystems.

Resistant individuals can be identified putatively via field assessments (Makinson, 2018) and experimentally through artificial inoculation with the pathogen (Sandhu & Park, 2013). These methods are effective but can be labour and resource intensive. Testing of individuals requires specialised facilities and takes weeks after seedling treatment to confirm disease resistance status following inoculation of 6-month-old seedlings. However, experimental testing may be avoided if it were possible to identify genetic markers linked to a resistance phenotype and use these to accurately predict resistance based on an inexpensive molecular assay (*e.g.*, Stocks et al., 2019; Weiss et al., 2020). This is the goal of genomic prediction models, which have previously been used successfully in both conservation (Fletcher, 2023; McGaugh et al., 2021) and commercial scenarios (Merrick et al., 2021; Ornella et al., 2012; Tan et al., 2017; Tsai et al., 2020).

Our study focuses on the foundation tree species *Melaleuca quinquenervia* (Cav.) S.T. Blake. This species exhibits heritable variation in myrtle rust resistance (Pegg et al., 2018) and is actively used in restoration of swamp forest vegetation. We aimed to support wetland restoration using a combination of experimental phenotyping, whole genome sequencing and association studies to identify predictive marker sets. Our goal was to identify genomic regions and variants that putatively facilitate host resistance to myrtle rust and use these to develop a rapid and low-cost method to predict resistance. We successfully achieved these goals by performing a Genome Wide Association Study (GWAS, *N* = 492 samples) for resistance to myrtle rust in *M*. *quinquenervia*. This generated a panel of associated markers used in the development of a genomic prediction model. Overall, the success of this study in detecting key genomic regions associated with myrtle rust resistance and in developing models with substantial predictive power provides a useful framework for other species that are impacted by myrtle rust or other pathogens. In summary, this study will enable us to work with on-ground practitioners to develop strategies for restoring populations that are enriched for disease resistance.

## Material and Methods

### Study species

*Melaleuca quinquenervia* is a dominant tree in many wetlands across the east coast of Australia. It is an ecologically important foundation species that helps maintain water quality through nutrient uptake, serves as an internationally recognised migratory bird habitat, and as an important food source for nectivores (Benson & McDougall, 1998; Bolton & Greenway, 1999; Grover & Slater, 1994). As such, significant impacts on this species would have much broader ecosystem consequences (also seen in Stevenson et al., 2023). High intraspecific variation in resistance to myrtle rust has been previously observed in *M. quinquenervia*, with variation both within and between tested populations (Pegg et al., 2018; Rayamajhi et al., 2010; Sandhu & Park, 2013). This intraspecific variation has been observed to be heritable (Pegg et al., 2018), suggesting that *M. quinquenervia* is suitable for a study of the genetic basis of resistance.

### Sampling

Leaf tissue samples and seeds were collected from 197 trees at 16 sites across the range of *M. quinquenervia* in the state of New South Wales (NSW), Australia (Fig. 1). GWAS was conducted over this extent due to limited genetic differentiation across this range (Fig. S3), and to reduce the representation of regions where introgression between *M. quinquenervia* and related species is possible (especially *M. leucadendra* and *M. viridiflora*, Edwards et al., 2010, 2018) (Fig 1c). At each site we sampled 6 to 22 trees, with the aim to space trees at least 20 m apart.

**Figure 1.**
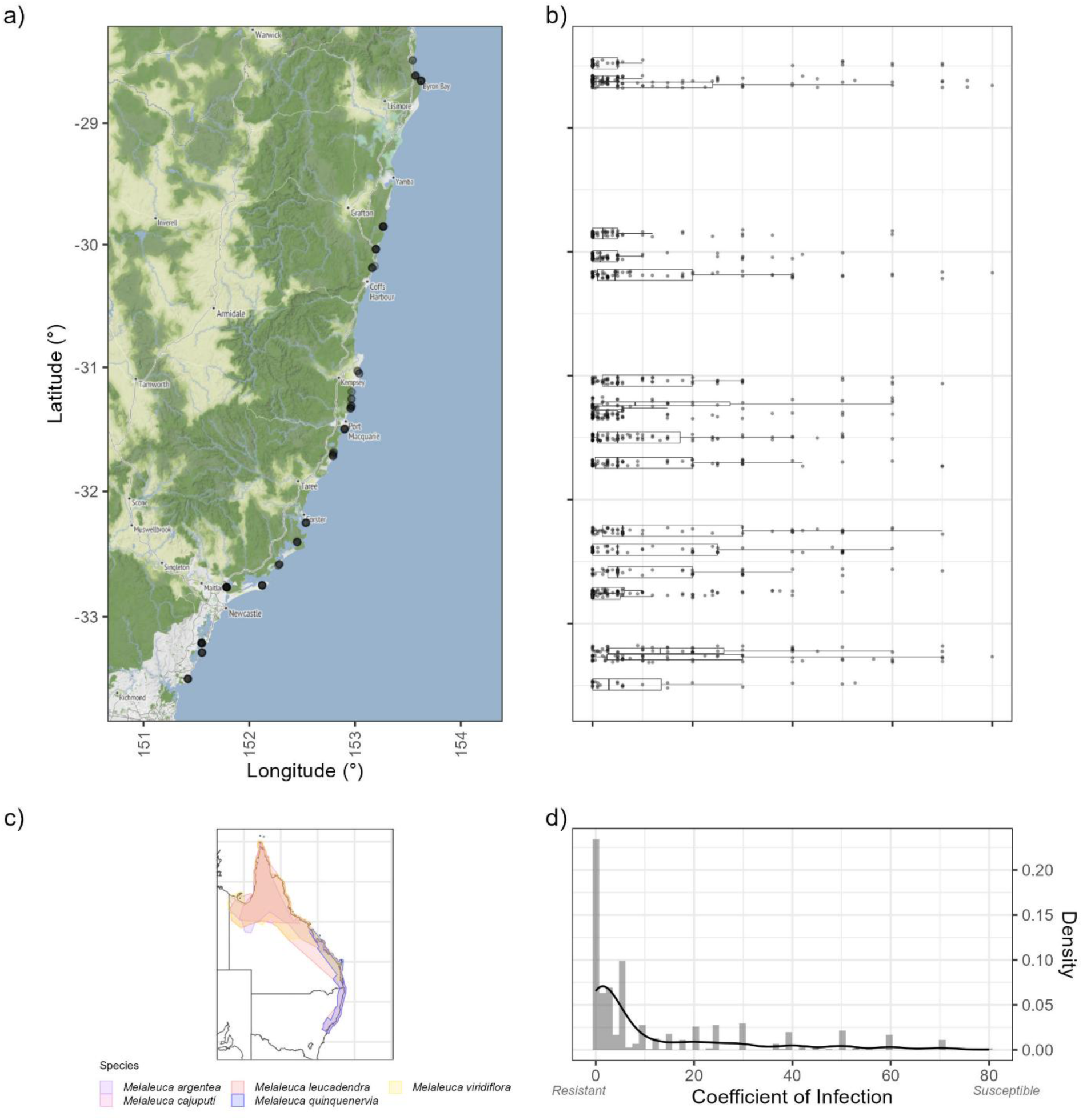
Maternal lines of Melaleuca quinquenervia sampled across NSW and the coefficient of infection (COI) of offspring. a) Each black dot represents a maternal line collected. b) Each box and jitter plot includes aggregated population groups at corresponding latitudes. Box plots represent the distribution of the coefficient of infection for seedlings of maternal lines within the aggregated populations. Overlayed jitter plots indicate seedlings and their coefficient of infection. c) Distribution of species from the broad-leaved Melaleuca clade across eastern Australia, with observations sourced from Atlas of Living Australia. d) Histogram and density frequency polygon of seedling COI distribution across all maternal lines.

To place our GWAS study in a broader evolutionary context, we obtained leaf tissue samples of *M. quinquenervia* from 20 samples across 4 sites in Queensland (QLD), and from representatives of several congeneric species. *Melaleuca quinquenervia* is part of a clade of 14 broad-leaved *Melaleuca* species where introgression is known to occur. We focused on the southern portion of the range for GWAS as in the north, ranges of closely related species overlap and introgression is potentially more common (Edwards et al., 2010, 2018). We also collected samples from four additional species from the broad-leaved *Melaleuca* clade: *M. leucadendra, M. viridiflora*, *M. argentea* and *M. cajuputi.* We sampled two more distantly related species as potential outgroups (*M. nesophila* and *M. nodosa*) (see ‘Data and script availability’).

### Pathogen inoculum

We used the ‘pandemic’ strain of *A. psidii* maintained at the Plant Breeding Institute (PBI), with isolate ID: Au_3, PBI culture no. 622 and accession 115012.

### Rust assays

For rust resistance assays, approximately 12 seedlings per sampled tree were grown in small trays containing growth medium (1:2 coir peat to sand, with 435:500:365:2000 fertiliser of dolomite, iron sulphite, micromax and Osmocote). After 31 days, approximately 12 seedlings per sampled tree were moved to forestry tubes (50 x 50 x 120 mm) containing a mix of High Growth AFP growing media (1:1 mixture of 0-8 mm pine bark, 2-5 mm pine bark, coir peat, coarse sand), topped with a layer of perlite and doused with anti-fungal solution (1:600 Agri-Fos® 600 to water). Seedlings were grown in randomised positions on benches and watered by misting from above. For the estimation of relative growth rate (RGR), plant height measurements were taken at 139 and 163 days after sowing. Seedlings at 174 days after sowing were then screened for myrtle rust resistance.

Myrtle rust resistance assays followed a set of procedures described previously by Sandhu and Park (2013) with modifications. Briefly, a *Syzygium jambos* plant was inoculated with urediniospores of *A. psidii*. Approximately 20 mg of fresh urediniospores were collected from the *S. jambos* plant and suspended in 10 mL of isopar-L paraffin oil. For the inoculation of *M. quinquenervia* seedlings, the rust suspension was atomised using an airbrush attached to a motorised compressor. After inoculation, urediniospores were left to settle for 5 minutes, then seedlings were moved to a dark incubation chamber for 24 hours at 20°C and misted constantly to maintain >95% relative humidity. Following incubation, the seedlings were moved to a microclimate room maintained at 22 ± 2°C for 14 days, prior to being scored for rust infection and host response, as described below. Seedlings that showed no evidence of infection were reinoculated and scored a second time, to ensure initial scores were robust and seedlings were challenged by the rust spores.

### Rust Scoring

Rust infection scoring followed a protocol described previously by Sandhu and Park (2013). Briefly, rust response was described based on infection types including the presence or absence of active spores, chlorosis, and/or necrosis. Active spores were designated a numerical score based on its coverage, while the latter responses were presence or absence scores. Data on the infection type and disease severity were then used to calculate a single numerical coefficient of infection (COI), based on the modified Cobb’s scale (Peterson et al., 1948) for each seedling (protocol found in Supplementary Information). In addition to this, following the numerical score of infection, a binary classification of susceptible (numerical score > 2) and resistant (numerical score 0–2) was generated.

### DNA extraction, library preparation, and sequencing

DNA was extracted from leaf tissue samples by the Australian Genomic Research Facility (AGRF, Melbourne, Australia), using 96-well Qiagen DNeasy plant mini kits. Libraries were generated for short read sequencing using an approach described by Jones et al. (2023). Briefly, this protocol uses dilute transposase fragments to insert common adaptor sequences followed by PCR with custom barcode pairs. This produces whole genome coverage with a substantial reduction in cost, though some repetitive sequences may result in variable sequencing depths. In total, 752 libraries were multiplexed in two batches, and each was run in a single S4 flow cell on an Illumina NovaSeq 6000 sequencing instrument at the Biomolecular Resource Facility, Australian National University.

### Data analysis

Read ends were trimmed to remove low quality bases (≤ 24 quality threshold in windows of 8 bp) and adaptor sequences, and overlapping paired end reads were merged, using AdapterRemoval (v2.3.2; Schubert et al., 2016). Reads were mapped to the *M. quinquenervia* reference genome (‘HapA’; Chen et al., 2023) using BWA (v0.7.17-r1188, Li & Durbin, 2010). BAM files were processed using SAMtools (v1.6, Danecek et al., 2021), variants were then called using the ‘mpileup’ function in BCFtools (v1.17, Danecek et al., 2021) and filtered using VCFtools (v0.1.16, Danecek et al., 2011).

We performed specific analyses using datasets that contained different subsets of samples and applied filters that were suited to the inferential methods that were used. This resulted in three main datasets. The first dataset was used to perform association analyses and began with genotype data from 520 *M. quinquenervia* seedlings (including 2 technical replicates). For this dataset, we filtered out sites with a mapping quality lower than 15, removed insertions and deletions, and removed sites that were not bi-allelic. We then applied further filtering thresholds including a minimum depth per site of 6 and a minimum genotyping quality of 20. Sites that were missing in more than 200 individuals were excluded. Samples that had a mean depth of coverage of less than 4 were also removed, resulting in a final 492 samples. For several downstream analyses (noted below), a minor allele frequency of 5% was applied (2.4M SNPs). BEAGLE (v5.4, Browning et al., 2018, 2021) was used to perform imputation prior to GWAS.

The second dataset initially consisted of samples of 188 maternal trees from NSW, and 18 samples from QLD, with the final sample counts subsequently selected for analysis post-filtering. This dataset was used for association analyses and for inferences concerning population genetic structure and demographic history. For association analyses, we began with the initial 188 genotyped maternal trees from NSW. Of these, 182 samples remained after filtering, which was performed in an identical manner to the seedling dataset above (except the mapping quality filter of 15), and with imputation performed using BEAGLE (Browning et al., 2018, 2021). A minor allele frequency filter of 5% was applied to this dataset for association analysis (5.9M SNPs). For the analyses of demographic history and population structure, the dataset was initiated with all samples from NSW and QLD. This was subjected to the same filters, however was not imputed. A final dataset of 181 NSW maternal samples and 17 QLD samples were used. For population structure analyses, we anticipated that far fewer SNPs were sufficient for robust inference. Therefore, we thinned the dataset to 1 SNP every 4,000 bp (58,460 SNPs), balancing statistical reliability with computational efficiency.

The final dataset was used to make inferences at deeper evolutionary time scales, especially concerning introgression. For these analyses, we generated a variant file that initially included samples from all the available outgroup species, as well as samples from QLD. For computational efficiency, we used subsets of the samples from NSW, which are described below. These variant datasets were filtered with thresholds that were slightly less stringent, reflecting their application at deeper evolutionary scales, where failure to identify heterozygous from low coverage sites is less problematic. For these data, we removed insertions and deletions, and sites that were not bi-allelic, and imposed a minimum depth per site of 4 and minimum genotyping quality of 20.

### Population genetic inference

We examined patterns of population genetic variation among *M. quinquenervia* samples using clustering methods to characterise population structure, as well as methods that infer demographic history (effective population size). First, a Principal Components Analysis (PCA) was performed on the dataset containing NSW and QLD samples of maternal lines. Prior to this analysis, additional minor allele frequency (5%) and site missingness (threshold of 50%) filters were applied, along with thinning for computational efficiency (1 SNP per 4,000 bp). The PCA was implemented with ‘glPca’ from the package adegenet in R (v2.1.10, Jombart, 2008). We made inferences about population structure using conStruct (10,000 iterations, up to 6 ancestral populations analyzed; Bradburd et al., 2018), and sparse non-negative matrix factorisation (sNMF, implemented in R package LEA with entropy set to true and 9 repetitions for each of the 6 *K*-values; v3.20.0, Frichot & François, 2015). These analyses were performed on the dataset of maternal lines thinned to 58,460 SNP sites. For these analyses, a population from QLD with only 2 samples was excluded due to small population size.

We inferred the demographic history of *M. quinquenervia* using SMC++ (v1.15.4, Terhorst et al., 2017). Two iterations of this analysis were conducted. The first included samples drawn from large regions, including all NSW maternal tree individuals and samples from QLD, respectively. The second set of analyses consisted of separate SMC++ runs for the maternal trees collected from each sampling site. For each SMC++ analysis, three distinguished lineages were selected using individuals from the target populations of the analysis. These analyses used a masking file for repeats identified in the genome annotation (Chen et al., 2023) and were conducted with parameters of mutation rate of 1 x 10^-8^ and a polarisation error of 0.5.

We next sought to determine whether there was evidence of introgression between *M. quinquenervia* and other broad-leaved *Melaleuca* species, particularly in regions where it occurs in sympatry with closely related congeners. We conducted this by estimating the D statistic for a series of ABBA-BABA tests, which were implemented in Dsuite (v0.5 r48, Malinsky et al., 2021). Each of these tests asked if there was a difference, between two populations of *M. quinquenervia* (hypothetically, ‘P1’ and ‘P2’), in their tendency to share derived alleles with a closely related species of *Melaleuca* (hypothetically, ‘P3’). For all tests we used *M. nodosa* as the outgroup (to identify derived alleles). The tests considered possible gene flow between *M. quinquenervia* populations and closely related species including *M. cajuputi, M. leucadendra, M. viridiflora,* and *M. argentea,* which were represented by a single resequenced sample in their respective ABBA-BABA tests. All these species exist in the north of QLD, and the ranges of *M. viridiflora* and *M. leucadendra* extend into southern QLD (Fig. 1c). For *M. quinquenervia*, we divided the available samples into three populations. The first included *M. quinquenervia* samples from QLD, which are (broadly) sympatric with the four closely related species. The second included *M. quinquenervia* samples from northern NSW (north of −29.0 degrees latitude), which only are in proximity with *M. viridiflora* and *M. leucadendra* populations, relative to *M. quinquenervia* from the rest NSW. The third population included *M. quinquenervia* from the rest of NSW, which might be hypothesized to share disproportionately fewer alleles with the closely related species from the north. To reduce the computational demands of these analyses, we randomly selected 10 samples from each of the nominated NSW regions prior to performing the ABBA-BABA analyses.

Finally, we examined linkage disequilibrium (LD) decay across the genome to provide context for understanding the results of association analyses (Neale & Kremer, 2011). We did this using the same dataset that was used in seedling association analyses, filtered to exclude loci with < 5% minor allele frequency. Linkage disequilibrium (*R*^2^ estimator) was conducted with VCFtools at a window size of 10,000 bp.

### Phylogenetic inference

We estimated ancestral relationships among broad-leaved *Melaleuca* samples beginning with the dataset containing the outgroup species described above. For this analysis, we wanted to use a subset of the samples of *M. quinquenervia* from NSW, for the sake of computational efficiency, and to focus on the evolutionary relationships among plants with contrasting levels of myrtle rust resistance. We therefore selected samples to represent different levels of susceptibility and resistance against myrtle rust, including the most resistant (breeding value of 4.15 to 5.63) and susceptible NSW maternal trees (breeding value of 21.78 to 27.80), based on estimated breeding values of COI (see Association Analyses, below).

For these samples, we used the variant file, in combination with the reference genome sequence, to generate aligned sequences for a set of target loci. We did this using samtools consensus for 2,304 putative single copy genes (based on BUSCO v5 annotations, Simão et al., 2015; Chen et al. 2023). We trimmed sites with excess missingness (> 50%) using ClipKIT (v2.3.0, Steenwyk et al., 2020). We then inferred a gene tree for each alignment with IQ-TREE2 (v2.2.2.7 Kalyaanamoorthy et al., 2017; Minh et al., 2020a, 2020b), using ModelFinder to determine the best-fit model and estimated 1,000 ultrafast bootstrap replicates and branch tests (SH-aLRT) with 1,000 replicates (Hoang et al., 2018). A summary coalescent tree was inferred with ASTRAL (Mirarab et al., 2014) using all gene trees that were estimated using alignments with greater than 20 parsimony informative sites (i.e., 548 gene trees were dropped). Concordance factors were calculated at the site and gene level using IQ-TREE2 (gcf and sCFl using 100 quartets, Mo et al., 2023).

Finally, we examined the ancestral relationships among samples in genomic regions with a high density of sites associated with resistance to myrtle rust (see Results). To investigate this, we generated an alignment for a 200 kb region around the centre of the peak of associated SNPs on Chromosome 8 (positions 19,157,911 to 19,557,911) and estimated a gene tree for this region using IQ-TREE2 (4,955 parsimony-informative sites), implemented with the approach described above.

### Association analyses

GWAS analyses were performed using the statistical model GEMMA (v0.98.3, Zhou & Stephens, 2012) with a linear mixed model and a Wald test method. These analyses used a kinship matrix that was also estimated by GEMMA. In tests of significance, the Bonferroni correction threshold for multiple tests was set to 1 x 10^-7^. Multiple GWAS analyses were conducted using different parameters and approaches to ensure results were not sensitive to changes in these factors (Table S4). This involved changes in dataset filtering, the coding of phenotypes, and the use of different inferential methods. A final analysis is presented in the main text, and results of additional analyses are presented in Supplementary Material to illustrate the robustness of results across analysis settings.

Association analyses were performed for the COI phenotype that was measured for 492 seedlings (see above). COI values were transformed (4th root) to better meet normality assumptions for statistical testing (Fig. 1d, Fig. S6b). We also performed association studies for rust response phenotypes including a binary transformed infection score (Yong et al., 2021), and the rust responses of ‘flecking,’ ‘necrosis’ and ‘chlorosis’. Additionally, we wanted to check that markers of rust resistance were not associated with a simple growth trait, in a way that might have consequences for downstream use in restoration. We therefore also performed an association study for Relative Growth Rate (RGR). For maternal trees, breeding values were estimated using their seedling phenotypes. Breeding values characterise phenotype values for an individual based on a sampling of observed values for related individuals. Here we inferred a pedigree based on known maternal line information and treated paternity for the open-pollinated seedlings as unknown. Breeding values were then estimated using a best linear unbiased prediction (BLUP) with the function ‘remlf90’ from the R package breedR (v0.12-5, Muñoz & Sanchez, 2023). This provided breeding values of maternal lines of COI and RGR for the GWAS, and additionally an estimate for the heritability of COI. Mean COI values from the seedling dataset were also tested as an additional phenotype. Both seedling and maternal genotype datasets were filtered as above, along with an additional filter of 5% minor allele frequency.

### Annotation of SNPs that were associated with resistance

To assess the impact of the associated SNPs we investigated their modifications of amino acid sequences. We used SNPeff (v5.2, Cingolani et al., 2012) to annotate 500 SNPs that were strongly associated with rust resistance (COI). We observed that a set of highly associated SNPs occurred near a group of NLR or NBARC genes on Chromosome 8 (Chen et al., 2023; see Results). We further examined the genomic variation in this region, in two ways.

First, for the small group of NLR genes in this region, we generated and compared haplotype sequences for each seedling. We did this using the (phased) variant file and the reference sequence, along with packages SAMtools (v1.6, Danecek et al., 2021) and consensus BCFtools (v1.17, Danecek et al., 2021).

Second, we examined variation in this region between haplotypes A and B of the phased reference genome (Chen et al., 2023). We did this after noticing that the tree used for the reference genome was heterozygous for several highly associated SNPs in this region. We first aligned Chromosome 8 from haplotypes A and B using AnchorWave (v1.2.5, Song et al., 2022), to find the region of haplotype B corresponding to this cluster of associated SNPs. For the genes in this region on each haplotype, we BLASTed sequences to the opposite haplotype. For further detail, please refer to the Supplementary Information ‘Gene annotations in a key region of Chromosome 8’. Putative matching genes were visualized using gggenomes (v1.0.1, Hackl et al., 2024) in this region.

### Genomic prediction and restoration scenarios

Next, we sought to develop Genomic Prediction models with the capacity to predict rust resistance (COI) phenotypes using genotype information at modest numbers of SNPs. We used SNPs identified by GWAS, and fit genomic prediction models to COI data using the method of genomic BLUP (gBLUP; Clark & Van Der Werf, 2013), implemented in the GAPIT R package (v3.5, Wang & Zhang, 2021). This approach produces a Genomic Estimated Breeding Values (GEBV) for each analysed sample.

Several iterations of the genomic prediction model were tested before finalising our model. We varied the number of SNPs, the size of the training and testing datasets, and applied different SNP filters (see Supplementary Information for a table of models and outcomes). Initial iterations of the models also included markers of higher density within clusters. Subsequent iterations of these models investigated the impact of the thinning of these clusters to reduce over-training of the model and the emphasis on major effect genes in exchange for minor effect genes. We also tested models that included panels of random SNPs chosen by thinning across the genome (Table S10, Iteration 24). When used alone (i.e., without associated SNPs), these provided a null expectation for the predictive capacity of models based on associated marker SNPs. However, we also anticipated these random SNPs would be included in panels that would be used for population management applications, where they could be used to infer relatedness and diversity of sampled individuals. Model performance was evaluated using metrics including model reliability (Wang & Zhang, 2021), and the Pearson correlation between predicted COI (GEBV) and the actual observed COI value for individual samples The evaluation metrics were calculated using 50 individuals that were chosen at random to be ‘held out’ from the training dataset. This meant we assessed model quality based on the observed and predicted COI values for individuals that were not used to train the model. For the final genomic prediction model, we used alternative methods for choosing individuals to be held out of analyses, as a check of the robustness of the model (Table S5–6). In particular, we checked that the model could make useful predictions for individuals from families or populations that were not specifically used in training. We held out individuals from randomly selected families (99–127 individuals), and from randomly selected populations (56–87 individuals). These analyses were repeated 5 times with different randomisation sets. We also specifically held out populations at the extremes of our geographical sampling to check that the predictive capacity of the model extended across the full sampled range (Table S9). Our final model was selected based on goals of restoration applications which require accurate predictions at minimal cost.

Previous studies have identified epistatic interactions between NLR genes that function in plant immunity (John et al., 2022; Phan et al., 2016). Our goal was to find candidate loci that might explain variation in COI via epistatic interactions, and to test if those loci could improve the performance of genomic prediction models. For these analyses, COI was converted to a binary ‘case/control’ phenotype where individuals with a COI score of 0 was designated as a Case, and remaining individuals as Control. ‘Fast-epistasis’ was first run with BOOST (gap of 1,000 kbp, Schüpbach et al., 2010, Wan et al., 2010) on the imputed seedling dataset, split by chromosome, to find a preliminary set of marker pairs that participated in epistatic interactions. We filtered this preliminary set of marker pairs, only keeping pairs that included a SNP site that exhibited an association with the phenotype (p-value < 0.001). For these SNPs, we performed a more in-depth analysis of epistasis using PLINK’s ‘epistasis’ function. We incorporated SNPs from this analysis into the final genomic prediction model by including a subset of 50 SNPs having the strongest epistatic interactions with SNPs from the original panel of associated sites.

## Results

### Rust resistance

*Melaleuca quinquenervia* seedlings varied substantially in their rust responses, from highly resistant to highly susceptible. This variation was observed both among and within families, and among and within populations (Fig. 1). Overall, based on the analyses of 1,098 seedlings across 183 families, the COI had a high level of heritability (*h^2^* = 0.87, estimated from the BLUP animal model), and inferred breeding values (BLUPs) varied substantially among families (range: 4.06 to 27.87). Within families, there was often substantial variation in COI values among seedlings, especially in families that were relatively susceptible (large COIs). There was no association between the COI values measured for seedlings and their estimated relative growth rates.

### Broad patterns of genetic variation in *Melaleuca quinquenervia*

*Melaleuca quinquenervia* exhibited high levels of genetic variation, with approximately 5.9M SNPs detected across the genome based on the 182 samples from wild populations in NSW. Across this focal area of this study, there was little evidence of discrete population genetic structure and a weak signature of isolation by distance (Fig. S3). Based on samples from large regions of NSW and QLD effective population sizes exceeding 1 x 10^5^ were inferred for *M. quinquenervia* for much of the last 450,000 generations, before putative recent contractions (Fig. S4). Linkage disequilibrium between SNP markers tended to decay rapidly as a function of the distance between SNPs, with *R*^2^ values falling to half their initial value for SNPs that were less than 1,000 bp apart (Fig. 2, half of maximum value at 711 bp apart).

**Figure 2.**
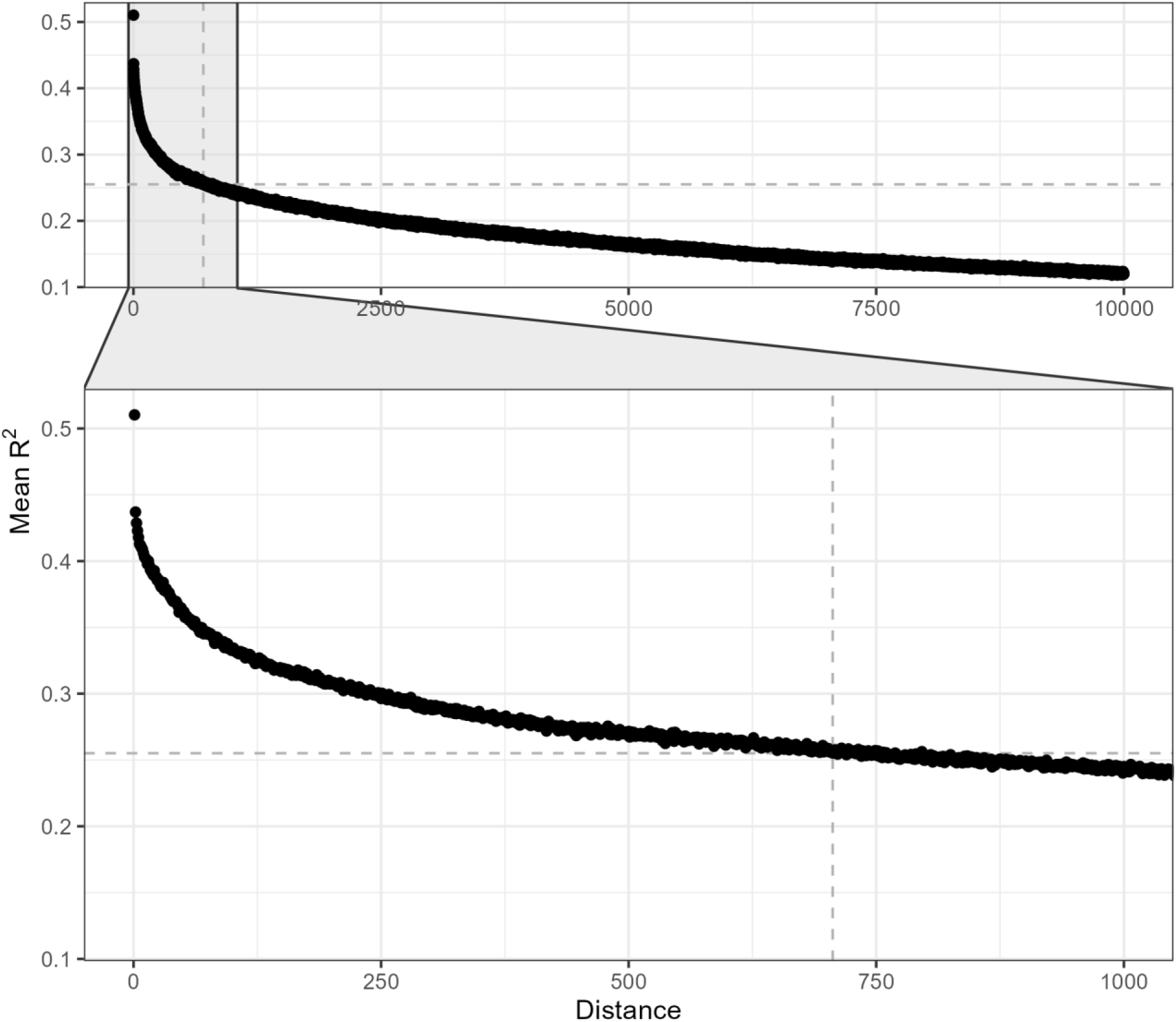
Linkage disequilibrium with increasing distance between SNPs across the whole genome, calculated with a window size of 10,000 bp. Markers represent mean R^2^ values aggregated by distance. The dashed line at distance 711 represents the R^2^ value of 0.254, representing the half of the maximum value.

The ABBA-BABA analyses, in aggregate, suggest northern populations of *M. quinquenervia* (QLD) had a greater tendency than southern populations (NSW) to share derived alleles with several closely related species. More specifically, *M. quinquenervia* populations from QLD showed significantly elevated sharing of derived alleles with *M. viridiflora* and *M. argentea* samples, relative to *M. quinquenervia* populations from northern or southern NSW (D statistics > 0.13, p-value < 1 x 10^-16^, Table S3, rows 1, 2, 4 and 5). However, *M. quinquenervia* populations from northern NSW did not have a disproportionate tendency to share derived alleles with these species relative to *M. quinquenervia* populations from southern NSW (Table S3, rows 3 and 6). For *M. cajuputi* the outcome was similar, but exaggerated, with ABBA-BABA placing *M. cajuputi* and *M. quinquenervia* from QLD as sister populations, but with disproportionately high frequencies of shared alleles between *M. quinquenervia* populations from QLD and NSW (Table S3, rows 7 and 8). Similarly, the sample of *M. leucadendra* was placed as sister to *M. quinquenervia* from southern and northern NSW, with *M. quinquenervia* from QLD sister to these (Table S3, rows 7 and 8).

### Genetic associations

Genome Wide Association Studies were performed between measures of COI and SNP genotype data. These analyses revealed a few major genomic regions containing SNPs that were clearly associated with COI (Fig. 3a), peaking above even a conservative Bonferroni adjusted threshold (p-value < 1 x 10^-7^). In particular, this included two regions on Chromosome 8, approximately 4 Mbp apart, containing 13 and 110 associated SNPs in each peak respectively, and a region on Chromosome 4 containing 15 associated SNPs. These associations are illustrated for several exemplar SNP loci in Fig. 3b.

**Figure 3.**
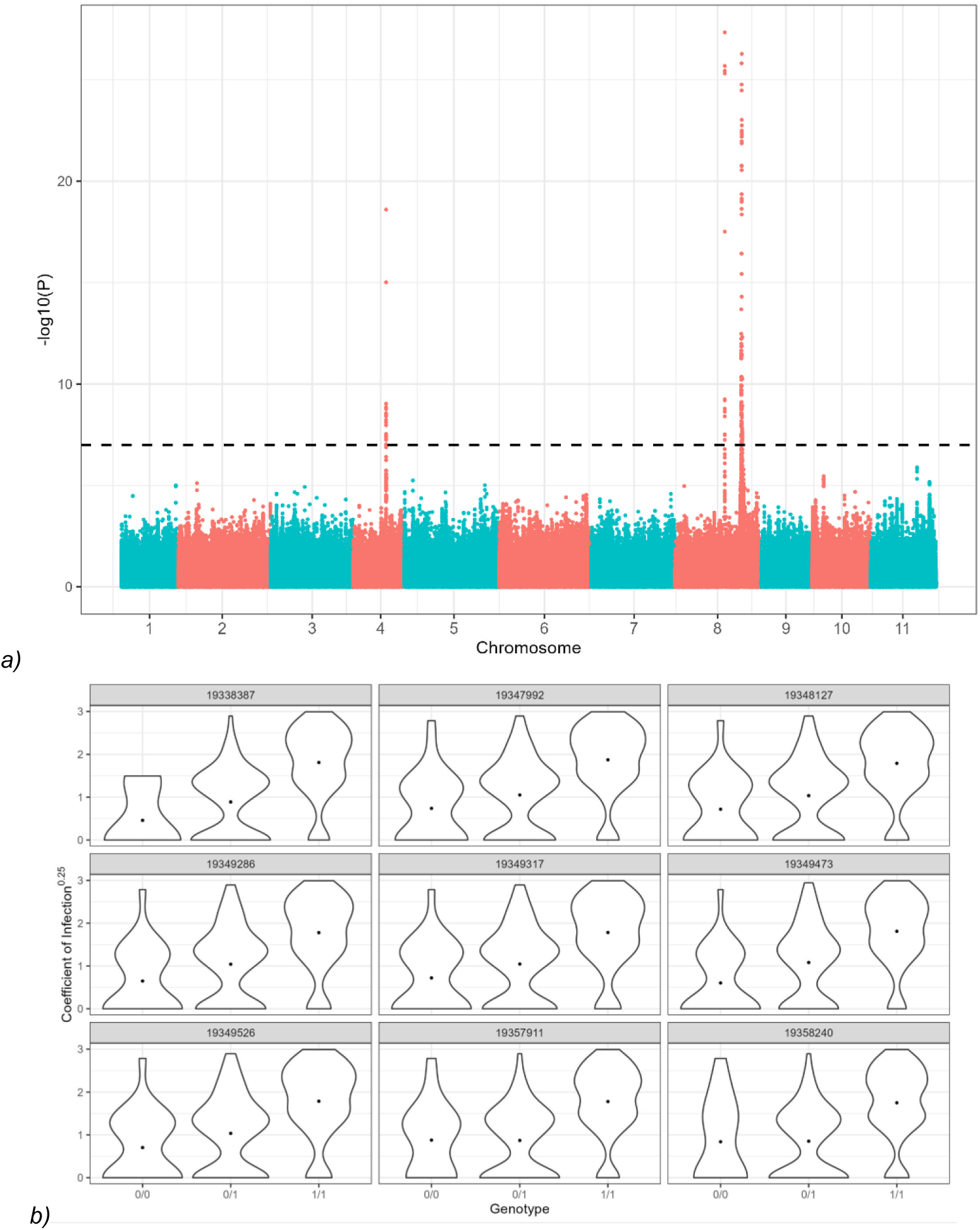
Genome Wide Association Analysis of the seedling dataset. a) Manhattan plot illustrating the strength of associations between SNPs along the Melaleuca quinquenervia genome and the coefficient of infection (COI; 4th root transformed). Three clear peaks of associations are observed. The horizontal dotted line marks the threshold of 1 x 10^-7^. b) Violin plots for a selection of significantly associated SNPs on Chromosome 8. These plots represent the spread of COI values for individuals with different genotypes at the nominated locus.

Associations in these regions were robust across a range of different analytical approaches and across different ways of encoding and measuring infection phenotypes (Fig. S6). For instance, when SNPs were filtered with greater stringency, and when we tested associations using different classifications of seedling infection, we observed significant associations in the same regions on Chromosomes 8 and 4 (Fig. S6, Table S4). Associations were also tested for different infection types and phenotypes. The phenotype ‘flecking’ was associated with SNPs in regions similar to those for COI, on Chromosome 8 and 4. However, association analyses of phenotypes ‘necrosis’ and ‘chlorosis’ produced none or only weak associations, distributed throughout the genome (Fig. S7).

A total of 2,100 SNPs that were associated with COI (p-value threshold of 1 x 10^-3^ and filtered with a Hardy-Weinberg equilibrium p-value of 1 x 10^-5^) were screened for epistatic effects on COI with SNPs from the rest of the genome. Many interactions were detected among the 2,100 associated SNPs, with 1,674 of them interacting epistatically with each other in association with resistance (asymptotic p-value < 0.05). Beyond this, another 426 SNPs participated in epistatic effects on COI. Of these 426 SNPs, 11.2% were on Chromosome 8. This was followed by 10.6% on Chromosome 7, 10.3% on Chromosome 5, and 9.86% on Chromosome 11.

A GWAS was also performed using the maternal dataset (*N* = 182). We tested associations between maternal genotypes (5.9M SNPs) and BLUP breeding values for COI. This resulted in several significantly associated SNPs distributed sparsely across the chromosomes (Fig. S8a) and included some associations that overlapped with regions on Chromosome 8 that had densely distributed associated SNPs in the analysis of the seedling dataset. We confirmed that these results were largely similar if we instead used mean seedling values as a measure of COI for a maternal tree, rather than BLUP breeding values.

In light of potential downstream ramifications in restoration, we wanted to confirm that loci associated with disease resistance were not associated with simple growth trait. We therefore tested associations between SNP genotypes and the phenotype Relative Growth Rate (RGR). When tested using our seedling data, we did not observe any significantly associated SNP sites. We also tested these associations at the level of maternal trees using estimated breeding values and again did not observe significant associations (Fig. S8c). These results suggest this growth trait is independent from genetic control of the myrtle rust disease.

### Functional annotation of SNPs that were associated with infection and immunity

Broadly, several genomic regions containing many SNPs that were associated with resistance phenotypes were annotated as containing NLR genes in the *M. quinquenervia* reference genome (Fig. 4). We investigated the annotations of highly associated SNPs using SNPeff (Table S2). Results from this analysis showed that the associated SNPs occurred near or within a suite of different genes, including Gene 97 (abbreviated g97, ID from Chen et al., 2023), which was annotated as an NLR. The coding DNA sequence of Gene 97 shows nucleotide BLAST homology to the disease resistance protein (RPV1-like) in *Eucalyptus grandis* (NCBI Reference Sequence: XM_039300131.1; BLAST score of 1888, 79.81% identity over a query cover of 69%). When we further investigated the top 20 SNPs associated with disease resistance from our GWAS, we noted that they were concentrated around a small set of genes (g93, g97, and g98) on Chromosome 8. A number of the SNPs in this region exhibited an excess of heterozygosity, relative to Hardy Weinberg proportions, implying unknown gene duplications and this prompted a more detailed examination of the haplotypes of the reference genome in this region. In the reference genome, which was heterozygous at a number of associated SNP markers, there were several cases where one gene from Haplotype A was similar to multiple genes on Haplotype B, based on an AnchorWave alignment that established the syntenic region, and BLASTn and BLASTp alignments between the haplotypes (see Fig. 5). Interestingly, transposable element genes were also found between the annotated NLR genes on Haplotype A with no matching genes on Haplotype B. (A detailed analysis of inferred sequence variation in Gene 97 is presented in the Supplementary Information.)

**Figure 4.**
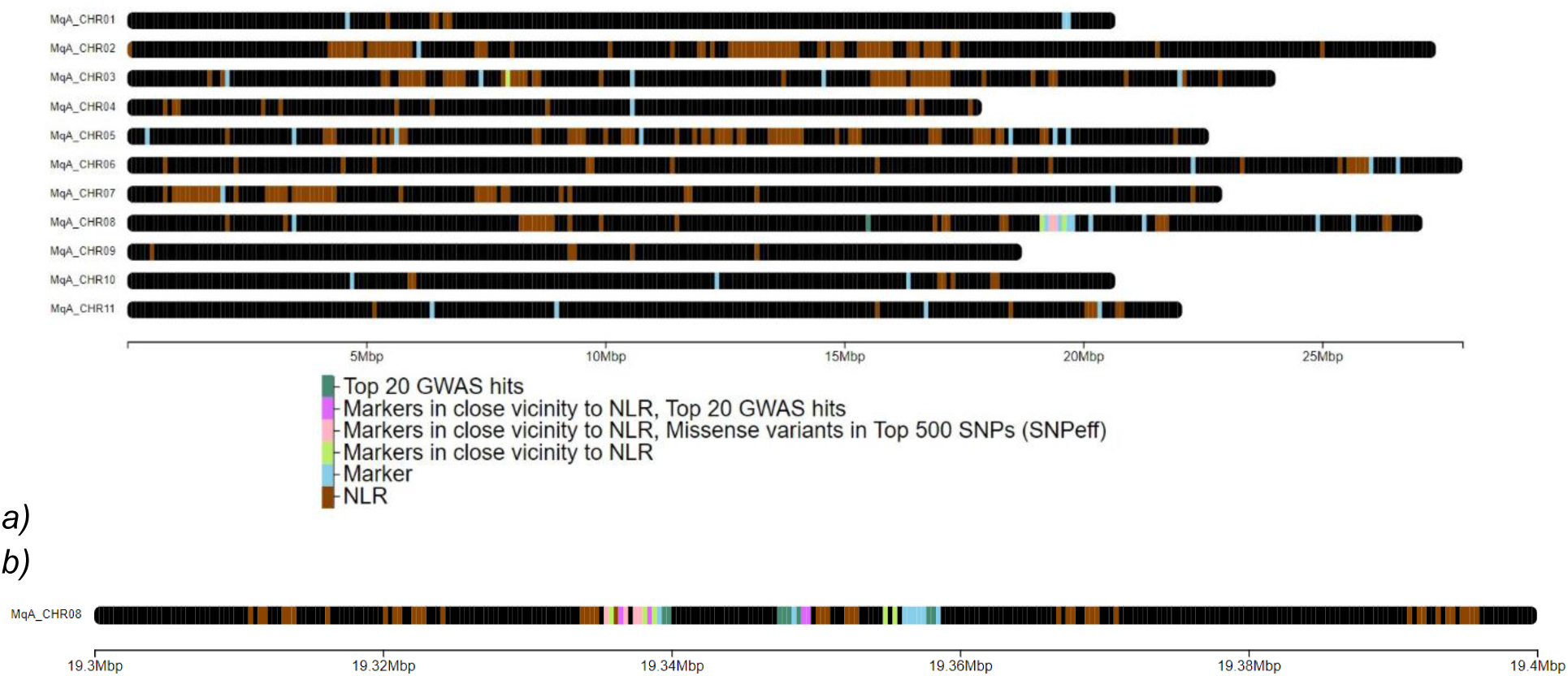
Chromosomes of Melaleuca quinquenervia and significantly associated marker SNPs from the seedling data genome-wide association study against coefficient of infection. Visualised using ChromoMap (Anand & Rodriguez Lopez, 2022). Brown markers represent the NLR regions annotated from Chen et al. (2023). Other markers represent coefficient of infection associated SNPs with a p-value lower than 5 x 10^-5^ coloured by their property. Markers are designated to be in close vicinity to NLR genes, specifically within a 1 kbp range. a) Whole annotated Haplotype A genome, b) Close up of the region associated with myrtle rust resistance (19,300,000 bp to 19,400,000 bp on Chromosome 8).

**Figure 5.**
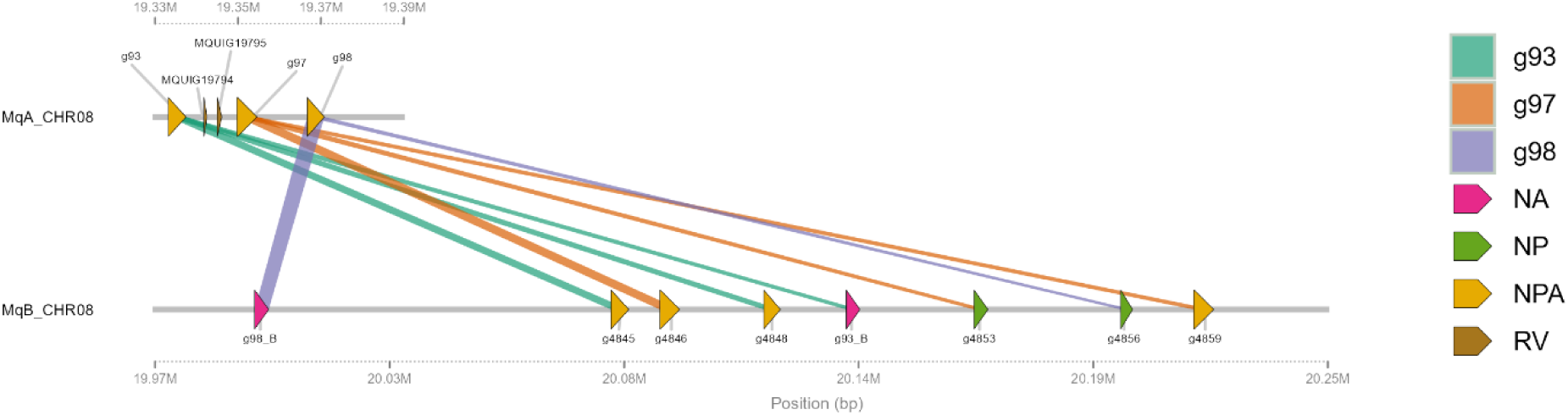
Genes g93, g97, g98 (ID from Chen et al. 2023) on Chromosome 8 of Melaleuca quinquenervia Haplotype A (sequence ID MqA_CHR08) and their matches on Haplotype B (sequence ID MqB_CHR08). The x-axis indicates base pair positions relative to the start of the sequence, with Haplotype A spanning from 19.33 Mbp to 19.39 Mbp, and Haplotype B spanning from 19.97 Mbp to 20.25 Mbp (ranges inferred from AnchorWave alignment). Links are coloured by their associated genes. Genes are coloured by their support, with ‘N’ corresponding to nucleotide BLAST support, ‘P’ corresponding to protein BLAST support, ‘A’ corresponding to AnchorWave alignment support. ‘RV’ is reserved for retrovirus transposons also annotated on Haplotype A. If a matching gene on Haplotype B is found for the corresponding gene on Haplotype A through protein BLAST results, they are labelled as such, otherwise they are labelled by the corresponding gene with a suffix ‘_B’. For more method details please refer to the Supplementary Information ‘Gene annotations in a key region of Chromosome 8’.

### Genomic prediction

A series of genomic prediction models were fit, leading to the identification of models and SNP sets that were strongly predictive for the COI phenotype, based on the criterion of performance in cross validation (Fig. S10). The process produced a ‘final’ model that had 1,049 SNPs and a cross-validation Pearson’s correlation of *R* = 0.83 (p-value < 0.01; Fig. 6). Several analyses confirmed that the cross-validation results for the final model were robust to different ways of choosing the individuals to be held out of analyses (Table S5–6). The 1,049 SNPs in the final model included 999 SNPs that had strong associations with the COI phenotype in the seedling GWAS. These 999 SNPs were part of a larger set of strongly associated SNPs, but the genomic prediction model improved after the SNPs were thinned according to their genomic location, by keeping 1 associated SNP per 1 kb window (Table S10). The remaining 50 SNPs in the final model were included because they took part in epistatic interactions that explained variation in resistance phenotypes, and because their inclusion in the model led to a modest improvement in prediction accuracy (Table S11, iteration 43).

**Figure 6.**
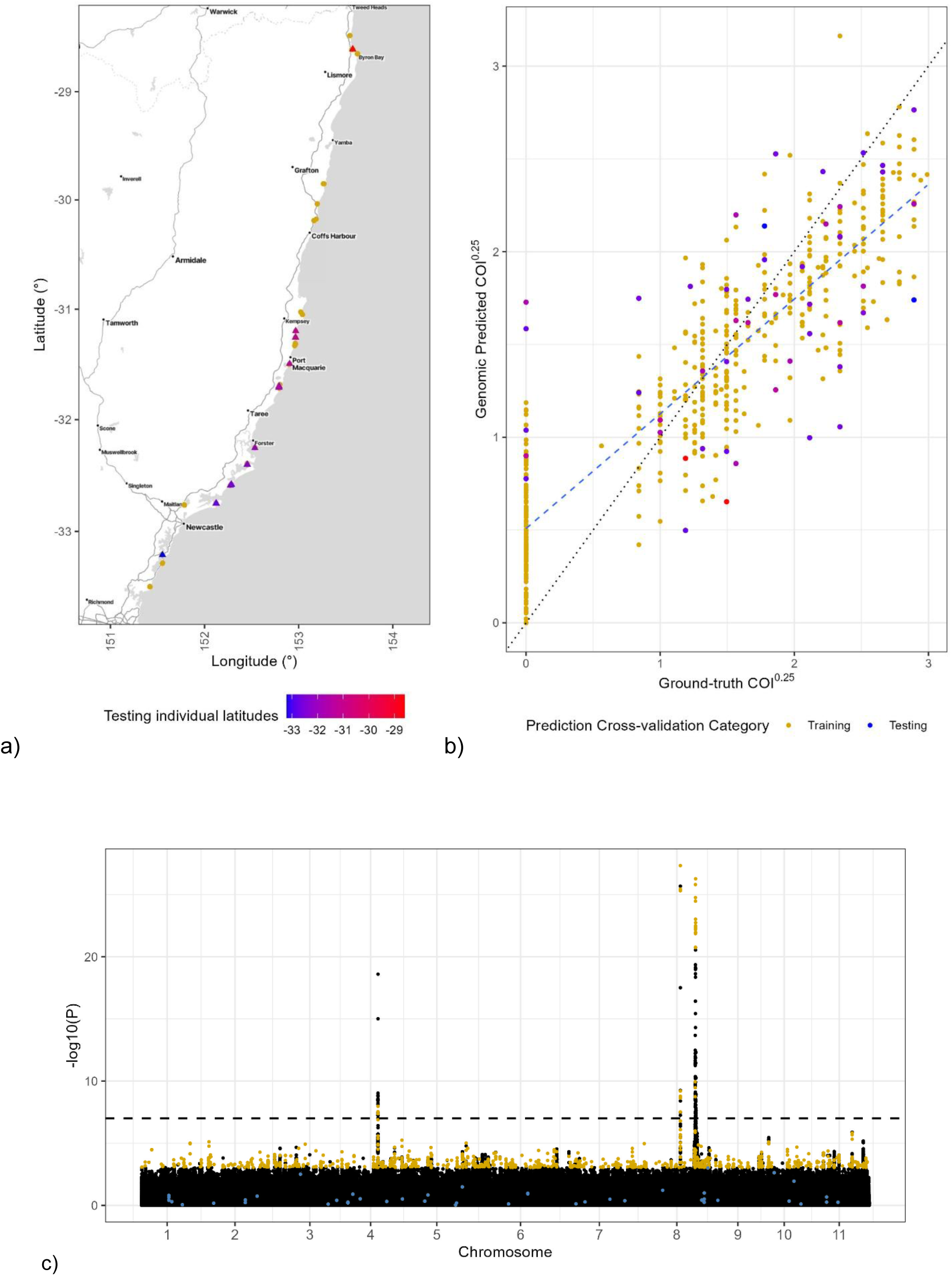
Genomic prediction model and the associated SNPs in the seedling GWAS Manhattan plot for COI. a) Points represent locations of the maternal lines collected for the seedling individuals in the genomic prediction model. Yellow points represent training individuals (492 individuals). The remaining points are used for testing (50 randomly selected individuals) and are coloured by the latitude of their maternal line. These individuals correspond to plot b). b) Plot of the predicted coefficient of infection from the genomic prediction model against ground-truth coefficient of infection. Each point corresponds to an individual and are coloured in accordance with the legend used in plot a). c) Manhattan plot illustrating the SNPs used in the genomic prediction model. Yellow points correspond to the resistance associated SNPs used in the genomic prediction model. Blue points correspond to the epistatic SNPs used in the genomic prediction model.

The predictive capacity of the final genomic prediction model was robust to the use of different ways of selecting the individuals to be ‘held out’ for model testing. Across different schemes for choosing individuals, Pearson’s correlation, *R*, between expected and observed COI values ranged from 0.79 to 0.85 (Table S5–6). These values were not affected by the exclusion of families or maternal lines but were associated with the number of training individuals used in the iteration, also explored in Table S9.

### Phylogenomic inference

At the whole genome-level, we estimated a species tree that generally placed *M. quinquenervia* samples together in a clade (Fig. 7). The other broad-leaved *Melaleuca* species were placed outside this clade, except for *M. leucadendra*, which was nested among *M. quinquenervia* samples from NSW (consistent with ABBA-BABA; Table S3). There was little suggestion that rust resistant or susceptible *M. quinquenervia* samples clustered together in this species tree.

**Figure 7.**
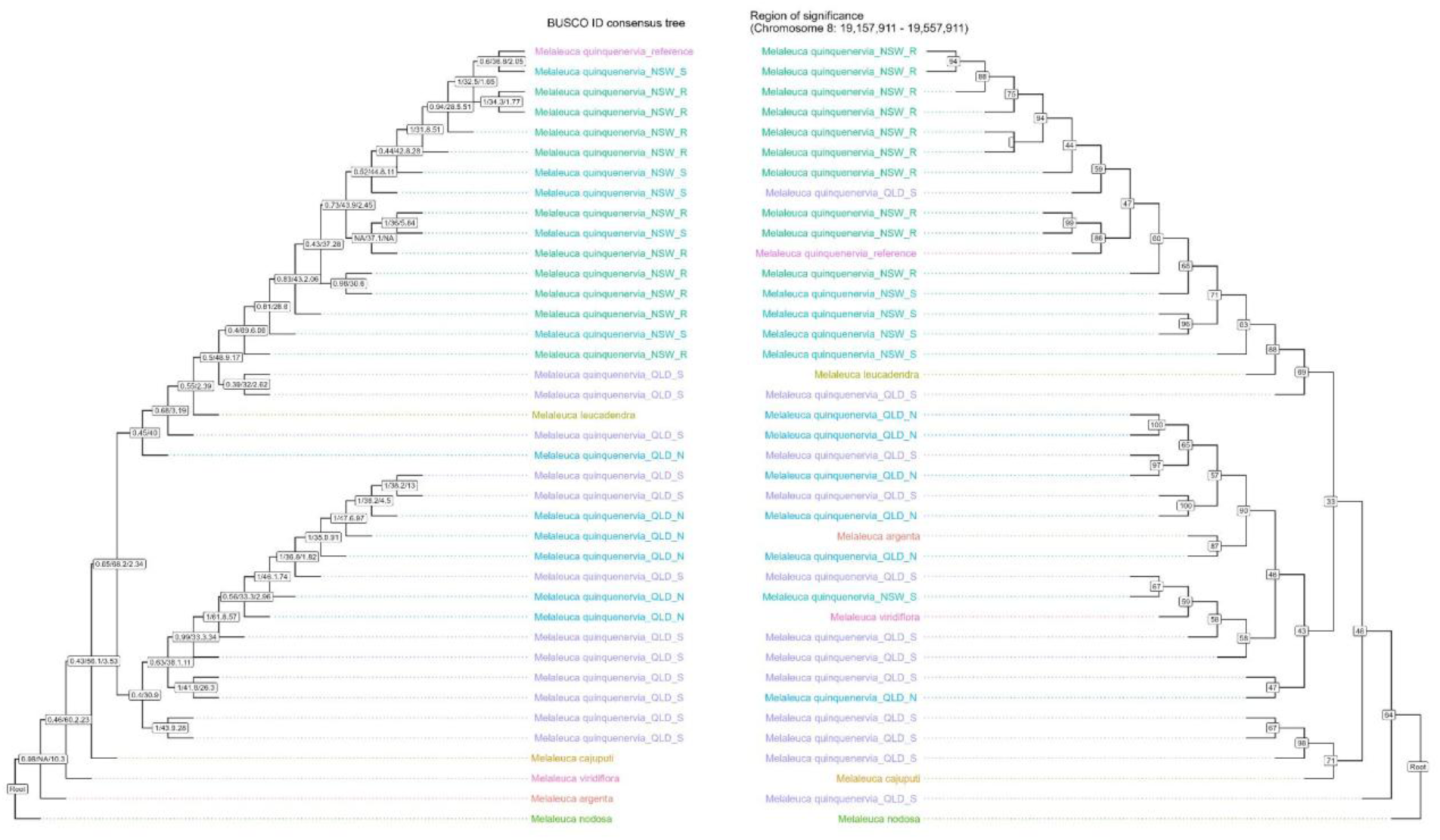
Phylogenetic trees estimated using (left) genome-wide BUSCO genes and a species tree approach and (right) a 200 kbp window flanking a region associated with myrtle rust resistance (19,157,911 bp to 19,557,911 bp on Chromosome 8). Tips include individuals from different broad-leaved Melaleuca species. Maternal lines in NSW are labelled according to myrtle rust resistance (R=resistant; S=susceptible) and QLD maternal lines are labelled according to geographical location (N=north; S=south). On the left, node labels of nodes correspond to ASTRAL multi-locus bootstrap support values/ sCFl / gCF generated on the final tree. On the right, node labels correspond to ultra-fast bootstrap support values.

We also estimated a gene tree for a region with many SNPs that were associated with myrtle rust (19,157,911 bp to 19,557,911 bp on Chromosome 8, containing 61 associated SNPs above the threshold of 1 x 10^-7^ and 4,955 parsimony-informative sites). Here, in contrast, resistant samples of *M. quinquenervia* from NSW largely formed a group to the exclusion of susceptible NSW individuals, albeit with poor branch support. The remaining samples, including the QLD *M. quinquenervia* and other broad-leaved *Melaleuca* species, did not form clades based on their regions of origin, and the outgroup broad-leaved *Melaleuca* samples were placed among the QLD *M. quinquenervia* samples.

## Discussion

Myrtle rust is impacting Australian ecosystems, threatening species (Carnegie et al., 2016), and altering the function and composition of plant communities (Fernandez-Winzer et al., 2019; McCarthy et al., 2024). We set out to develop molecular tools for predicting rust resistance in the wetland foundation tree species *M. quinquenervia*, with the goal of informing marker assisted restoration. This would involve choosing plants for restored populations based on their genotypes, to promote both myrtle rust resistance and genetic diversity. To do this, GWAS was used to identify SNP markers for resistance to myrtle rust, and these were incorporated into highly accurate genomic prediction models for this phenotype. Model development involved choices that balanced objectives of model predictive accuracy and the anticipated costs of deployment in restoration.

### Loci Associated with Myrtle Rust Resistance

Three regions of the *M. quinquenervia* genome contained high densities of SNP loci that were associated with myrtle rust resistance. This included two regions (approximately 4 million bp apart) on Chromosome 8, and one region on Chromosome 4. All three regions were in the vicinity of annotated NLR genes, which often function in responses to pathogens (Chou et al., 2023). At a finer scale, some of the SNPs that were most strongly associated with resistance to myrtle rust were located within putative disease-resistance genes and putatively encode non-synonymous substitutions (Fig. 4, Table S2).

One genomic region on Chromosome 8 that featured a cluster of strong phenotypic associations also appeared to exhibit structural variation. That is, annotated NLR genes on one haplotype of the phased reference genome showed sequence identity to two NLR genes in a corresponding region of the alternate haplotype (Fig. 5). These NLR genes were interspersed with transposable element genes. Genomic regions containing NLR genes tend to be highly dynamic (Barragan and Weigel 2021), with duplication being hypothesised to enable NLRs to recognise a wide range of effector proteins, conferring resistance to diverse pathogens (Dolatabadian et al., 2022; Li et al., 2021; Van De Weyer et al., 2019). These observations raise the possibility that markers associated with myrtle rust resistance reflect copy number variation, rather than simple SNP variation (e.g. if reads from multiple loci mapped to a single reference locus, Mastretta-Yanes et al., 2014). We have therefore been cautious about drawing inferences about precise molecular mechanisms. However, this does not affect our central goals if the markers remain predictive and the genomic prediction model remains robust when genotyped using different platforms. Our analyses also suggested it was helpful to thin the associated SNPs used in the genomic prediction model, perhaps because it reduced the concentration of markers in the region containing the putatively duplicated sequences.

### Prediction of Myrtle Rust Resistance and Deployment in Restoration

The key goal of our study was to develop molecular tools for predicting rust resistance traits rapidly and inexpensively, for applications in ecosystem restoration. To do this, we aimed to achieve levels of cross-validation accuracy comparable to genomic prediction models used in agricultural applications (*R* = 0.6, McGaugh et al., 2021), while using a relatively modest number of markers to limit genotyping costs. To this end, we designed a panel of 1,049 SNP markers which were incorporated into a genomic prediction model that achieved a high level of cross validation accuracy. The performance of this final model was robust to different cross validation treatments (Table S5–6). Overall, the final model was selected to minimise the number of markers without compromising predictive capacity (Table S7, Fig. S10). Broadly we found that model performance, determined by Pearson’s correlation, had diminishing returns with more markers. Once performance saturated, predictive power improved when we thinned markers (Table. S7, Iteration 27–30). This may occur due to large numbers of linked SNPs contributing relatively little independent information but promoting overfitting and potentially introducing noise (Song et al., 2024; Tan et al., 2017; Zhu et al., 2023). We also observed an improvement in predictive performance after incorporating a small set of markers that were involved in putatively epistatic interactions that explained variation in rust resistance phenotypes. This improvement in the genomic prediction model was modest but potentially represents a useful way to find additional markers that could be useful in models that do not depend exclusively on additive associations.

For deployment in restoration, the genomic prediction tool could be used to predict resistance in seedlings, or seeds from a mother tree. Making predictions for maternal lines and selecting seeds accordingly might be less costly, but also less accurate, than directly genotyping individual tube stock plants. The optimal way to incorporate this tool therefore requires consideration of the budget available for a project, the objectives for restoration, and the desired intensity of selection for resistance (Andres et al., 2024; Williams et al., 2024). For instance, if many seedlings were being planted and the circumstances did not require intense selection for resistance, it might be useful to perform genomic predictions for maternal genotypes, and to ensure a good representation of resistant seed lots were used for plantings. On the other hand, high value plantings in areas with strong disease pressure might benefit from genotyping seedlings and performing restoration using a subset of resistant and genetically diverse individuals.

### The Stability and Durability of Resistance

To be useful in restoration or conservation, disease resistance needs to be stable across environmental conditions, and durable over multiple generations of the pathogen (O’Hara et al., 2021; Sniezko & Koch, 2017; Sniezko et al., 2020). The resistance traits measured here were assessed in seedlings that had been inoculated with the pathogen under controlled conditions that were highly conducive to infection. This is highly useful for understanding responses of diverse genotypes under controlled conditions but might not reflect the diverse and dynamic conditions in natural environments where we aim to promote resistance for conservation and restoration outcomes. We therefore suggest it would be useful to monitor infection and resistance in genotyped individuals that are planted in natural environments (Chen et al., 2025). This will help better refine our understanding of the predictive performance of the model under different abiotic conditions, and especially those that prevail in realistic restoration scenarios. It might also provide new insights into the modulation of resistance by environmental conditions (including G-by-E interactions). This would involve the monitoring of disease severity across broader climate conditions, and across seasons, which has been observed to be important in crop and forestry studies (Mapuranga et al., 2022; Miranda et al., 2013). Disease resistance also potentially varies over the lifetime of plants (Mapuranga et al., 2022), or during the development of tissues (Beresford et al., 2020; Yong et al., 2019). It would therefore be useful to test the predictions of the model on adult trees, during broader studies in varying environments. However, we also note that seedlings are a critical life history stage, and their survivorship might be more impacted by myrtle rust infection than adult trees. For the goal of promoting restoration success, it is likely useful that the genomic prediction model directly addresses infection traits that were measured in seedlings.

Over longer time scales, several factors will potentially influence the durability of *M. quinquenervia* resistance to the myrtle rust pathogen. The resistance observed here appears to be dominated by several major loci, including those observed on Chromosomes 8 and 4. These multiple ‘R gene’ loci might provide greater durability than resistance mediated by a single locus. For example, rust resistance mediated by a single locus was lost in *Eucalyptus* plantations in Brazil, when it was overcome by a new variant of the pathogen (Almeida et al., 2021). This highlights that for restoration and conservation, it will be important to promote resistance using multiple alleles, where possible, to reflect polygenic selection in natural populations (Metheringham et al., 2025) and maximise the capacity for populations to adapt to changes in the pathogen. We note that it will be important to remain cautious about the breakdown of resistance, especially given the ability of the pathogen to complete its sexual life cycle on Myrtaceae hosts (McTaggart et al., 2018) and the possibility of incursions of new international variants.

### Broader Implications

This study concentrated on disease resistance in *M. quinquenervia*, especially where it occurs in southeast Australia, to avoid potential complications of interspecific gene flow with other species of the broad-leaved *Melaleuca* complex. This species is part of a group of 13 species, which exist as canopy trees in swamp habitats across vast areas of eastern and northern Australia. They have an important role in ecosystems that collectively have great cultural and biodiversity value and are increasingly recognised for their contribution to soil carbon storage (Jeffrey et al., 2020; Tran & Dargusch, 2016). The management of processes impacting broad-leaved *Melaleuca* trees potentially have substantial ecosystem level ‘multiplier effects’ in terms of biodiversity and biogeochemical function. It would therefore be interesting to know whether markers for rust resistance identified in *M. quinquenervia* also have predictive value in other broad-leaved *Melaleuca* species, and especially in those already known to exhibit variation in resistance (Pegg et al., 2018). This is particularly pertinent to northern populations of *M. quinquenervia* that likely share alleles disproportionately (relative to *M. quinquenervia* populations from further south) with several other broad-leaved *Melaleuca* species (Fig. 1c, Table S3). This is consistent with introgression between *M. quinquenervia* and other species when in sympatry or at close proximity (Edwards et al., 2018). This suggests it is quite plausible that the species of broad-leaved *Melaleuca* clade would share alleles that mediate disease resistance, or other traits, and suggests it would be useful to examine the markers identified here in individuals of other broad-leaved *Melaleuca* species, such as *M*. *leucadendra*.

This project has resulted in the development of a workflow for generating molecular predictions of resistance to myrtle rust in a species that is ecologically highly important. We look forward to further trials of the approach in natural environments, under diverse and changing climate conditions. This will allow us to explore the efficacy and cost efficiency of the approach for achieving targeted levels of resistance under different circumstances. We hope it might then serve as a template for research aimed at developing tools for marker assisted restoration in other species that are being impacted by myrtle rust, and by other pathogens.

## Acknowledgements

This research was supported by an Australian Research Council Linkage Grant (LP18010072) and funding from Transport for New South Wales. We are grateful to Joel Cohen for sample collection, and to Marlien van der Merwe, Pat Lu-Irving, Sam Yap, and Kit King for assistance in the glasshouse. We thank Joel Cohen, Bob Makinson, Marlien van der Merwe, Brett Summerell and Geoff Pegg for valuable advice at different stages of this project.

