## Supplementary Information for "Genomic prediction of disease resistance provides a path to marker assisted restoration in a wetland foundation tree species"

#### Contents

**Data and script availability**

**Protocol for scoring**

*Table S1. Immune type response score and their multiplicative factor.*

*Equation 1. Using both the coverage score and the immune type score to calculate the coefficient of infection for a seedling*

*Figure S1. Image examples of varying immune responses and coverages.*

**Supplementary figures**

*Figure S2. BLUP breeding values of Coefficient of Infection for* Melaleuca quinquenervia *maternal trees.*

*Figure S3. Genetic structure of the populations of* Melaleuca quinquenervia *maternal lines collected across eastern Australia.*

*Figure S4. Inferred historical demography (effective population size) of* Melaleuca quinquenervia *populations.*

*Figure S5. Principal component analysis of the SNPs in the thinned and filtered genetic data of maternal trees of* Melaleuca quinquenervia *(NSW and QLD).*

*Figure S6. Genome Wide Association Analysis of the seedling genetic dataset using different representations of phenotypes, untransformed coefficient of infection (COI) measurements and a binary transformation of the myrtle rust scoring scheme.*

*Figure S7. Genome Wide Association Analysis of the seedling genetic dataset for different infection and immunity phenotypes, necrotic, flecking, and chlorosis response.*

*Figure S8. Genome Wide Association Analysis of the maternal genetic dataset between the phenotypes of best linear unbiased predictions of breeding values for coefficient of infection, square-root transformed mean seedling COI by family, and best linear unbiased predictions of breeding values for relative growth rate.*

*Table S3. Results of ABBA-BABA analyses across the whole genome.*

**Seedling COI GWAS with different SNP filters**

*Table S4. Table of iterations of analyses conducted with different filters on the seedling dataset to determine the sensitivity of significantly associated MR resistance loci.*

*Table S2. Top twenty associated SNPs from the genome wide association analysis, and their associated SNPeff analysis results sorted by putative impact. Twenty-five missense variations present in the top 500 associated SNPs have also been included.*

#### Gene annotations in a key region of Chromosome 8

*Figure S9. Heatmap of gene 97's distance matrix when comparing two individual's allelic series.*

#### Genomic prediction

*Table S5. Randomly selecting 35 maternal lines used for testing.*

*Table S6. Maternal lines were grouped by population in accordance to location sampled.*

*Figure S10. Number of SNPs and the $R^2$ value between predicted COI and the ground-truth COI for individuals that were not used to train the genomic prediction model.*

*Table S7. Genomic prediction iterations focusing on changing the number of SNPs used and varying the filter of the genetic data.*

*Table S8. Genomic prediction iterations focusing on the markers discovered in the GWAS of the maternal genetic dataset and its application on predicting seedling phenotypes.*

*Table S9. Genomic prediction iterations focusing on varying the source and volume of the testing/training datasets.*

*Table S10. Genomic prediction iterations focusing on the inclusion of neutral SNPs from thinning the genome.*

*Table S11. Genomic prediction iterations focusing on using SNPs from alternative sources, including a GWAS that used BSLMM and epistatically significant SNPs.*

#### Data and script availability

Sequencing data are available on NCBI under BioProject Accession PRJNA1312870. Scripts and data for the figures in this publication are available in a Zenodo repository at https://doi.org/10.5281/zenodo.16792108. We note that two technical replicate samples were sequenced and incorporated into the seedling genotype dataset. One remained in the filtered dataset that was used in analyses. To ensure this did not affect outcomes, a replicate was removed for GWAS analyses presented in the main text, though it remained in iterations of the Genome Wide Association Analysis presented in the Supplementary Information.

#### Protocol for scoring

Scoring is completed by giving each individual a numerical score representing the maximum percentage coverage affecting a leaf on the individual plant (0-100, with 100 being full coverage). A second score of the immune type response is then given to each plant. For images and further information, see Sandhu & Park, 2013. This score can be used to determine a multiplicative factor (Table S1), which is used in Eqn. 1, to calculate the Coefficient of Infection (COI) for an individual plant.

*Table S1. Immune type response score and their multiplicative factor.*

| Immune type response | Description | Multiplicative factor |
|---|---|---|
| HR | Highly resistant | 0 |
| HRR | Highly resistant to resistant | 0 |
| R | Resistant | 0.15 |
| MR | Moderately resistant | 0.30 |
| MRMS | Moderately resistant to moderately susceptible | 0.45 |
| MS | Moderately susceptible | 0.60 |
| MSS | Moderately susceptible to susceptible | 0.75 |
| S | Susceptible | 1.0 |

*Equation 1. Using both the coverage score and the immune type score to calculate the coefficient of infection for a seedling*

$$Coefficient\ of\ infection = (Coverage\ score) * (Immune\ type's\ multiplicative\ factor)$$

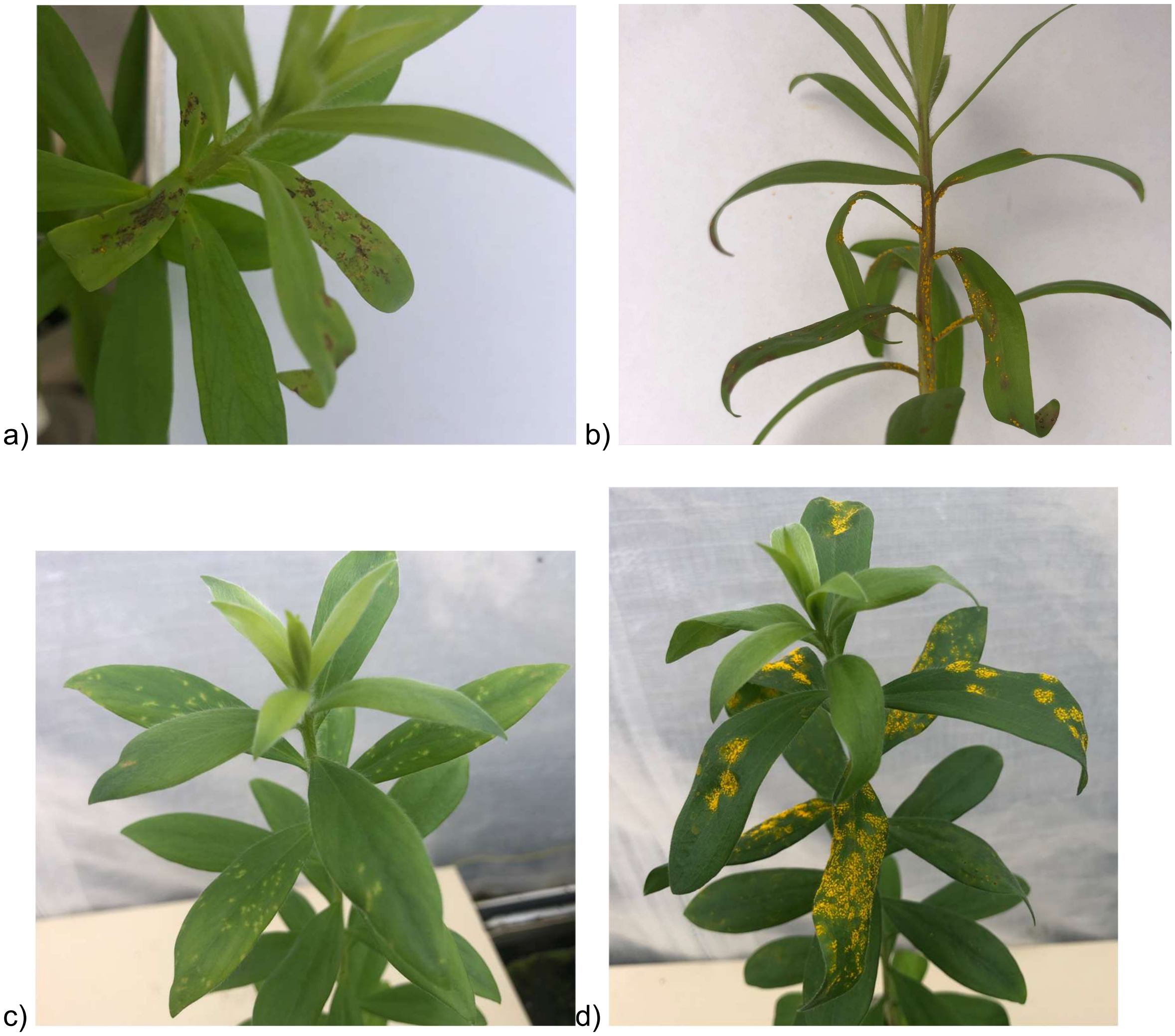


*Figure S1. Examples of varying immune responses and coverages. a) Individual with flecking, chlorotic, and necrotic spots. Rated as 30 coverage score and MR immune type, corresponding to a coefficient of infection (COI) of 9. b) Individual with necrosis, flecking and restricted sporulation. Rated as 30 coverage score and S immune type, corresponding to a COI of 30. c) Individual with flecking and chlorosis. Rated as 50 coverage score and R immune type, corresponding to a COI of 60. d) Individual with unrestricted disease sporulation. Rated as 60 coverage score and S immune type, corresponding to a COI of 60.*

### Supplementary figures

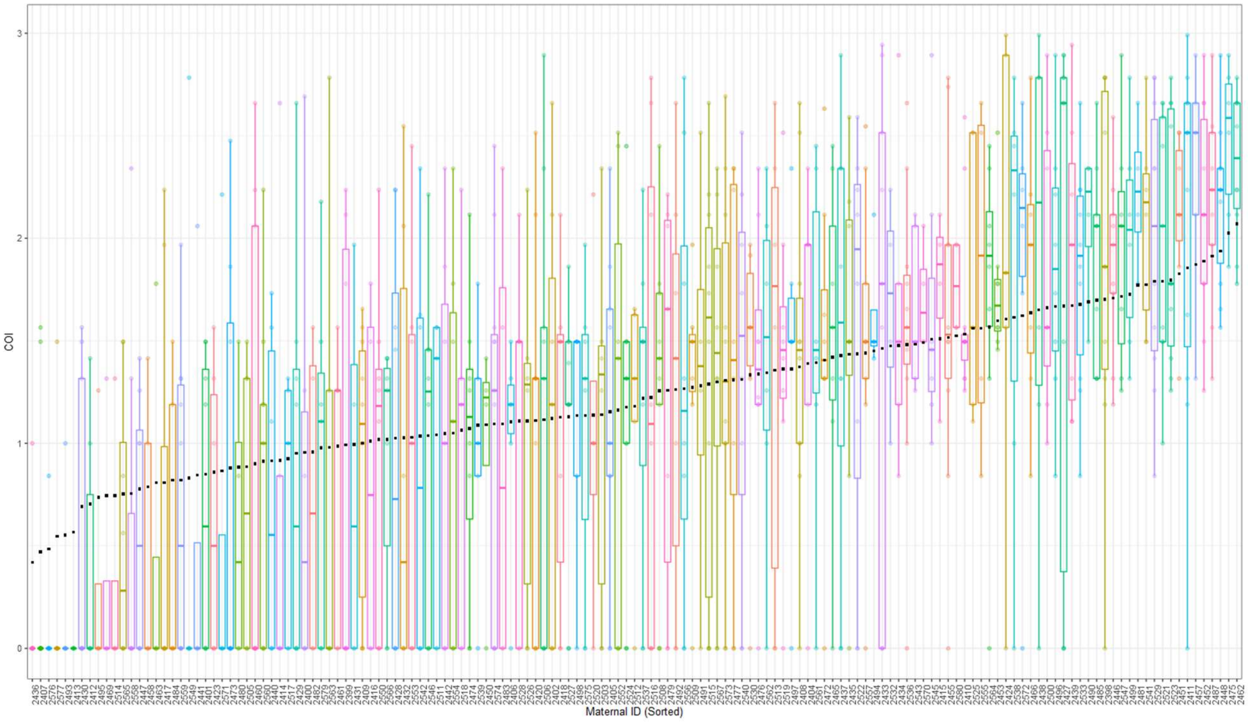


*Figure S2. BLUP breeding values of Coefficient of Infection for Melaleuca quinquenervia maternal trees. Each black square represents a COI for the BLUP breeding value of a maternal line, with the corresponding boxplot illustrating COI values of individual seedlings.*

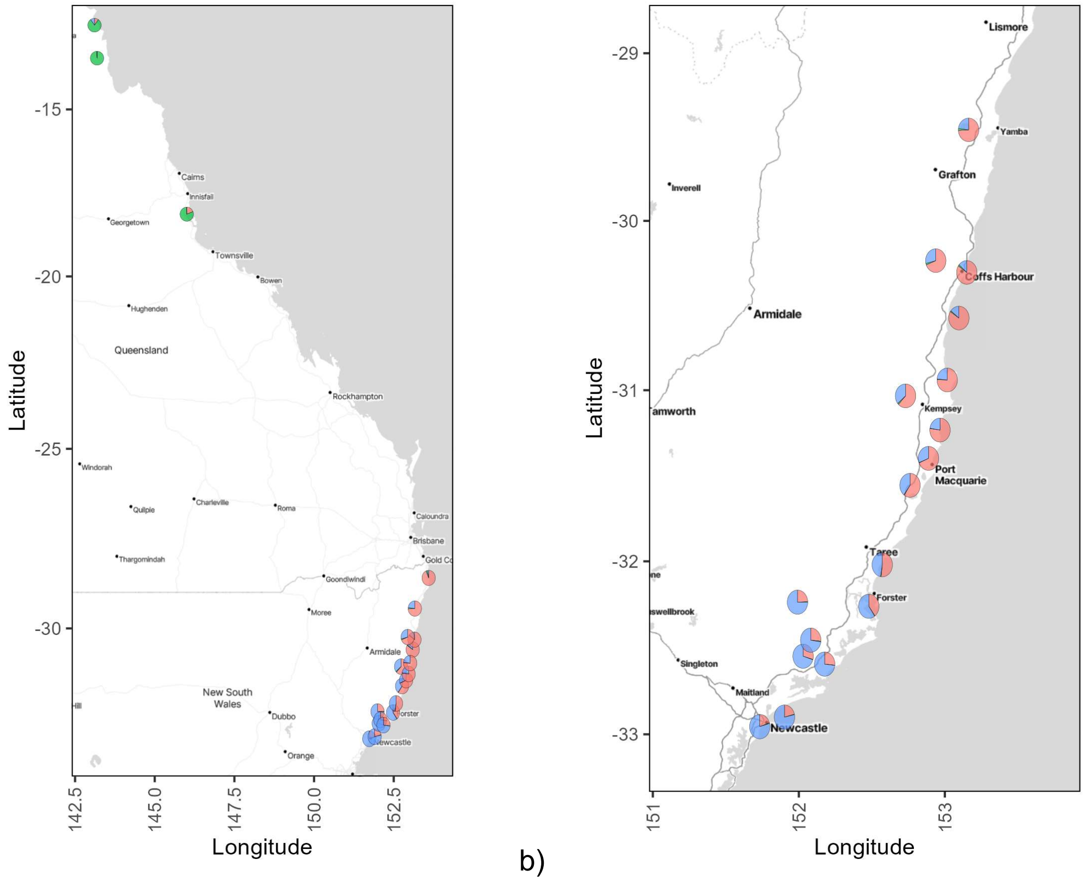


a) b)

*Figure S3. Genetic structure of the populations of* Melaleuca quinquenervia *maternal lines collected across eastern Australia. Pie charts for each population represent the mean ancestry coefficient values estimated by sNMF admixture analysis (Frichot et al., 2014; Frichot & François, 2015), modelled with K=3 ancestral populations. a) Both New South Wales and Queensland populations are illustrated. b) Only New South Wales* M. quinquenervia *populations are shown.*

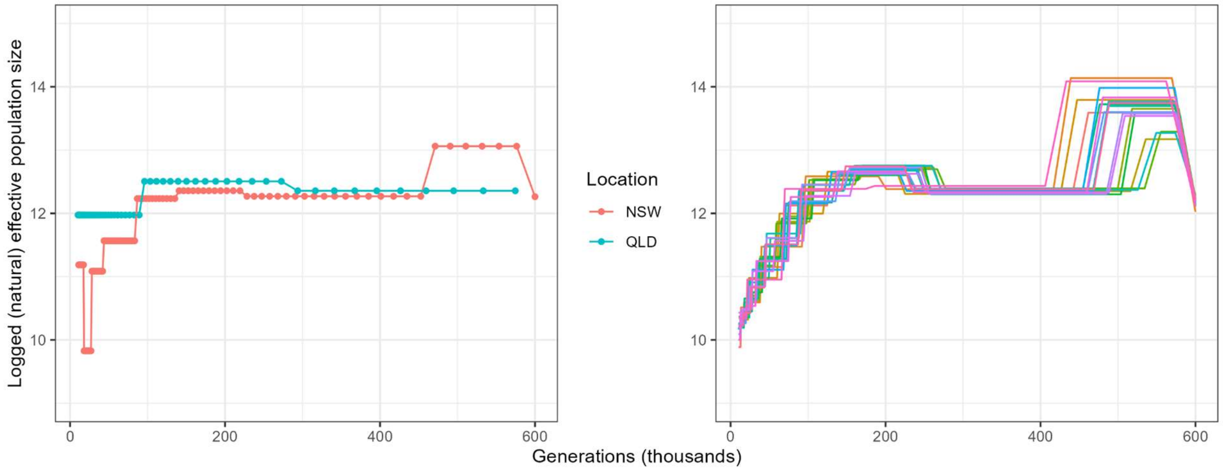


*Figure S4. Inferred historical demography (effective population size) of* Melaleuca quinquenervia *populations. The left panel separates the data by state, New South Wales and Queensland. The right panel further splits the samples by sites, designated from their collection locations.*

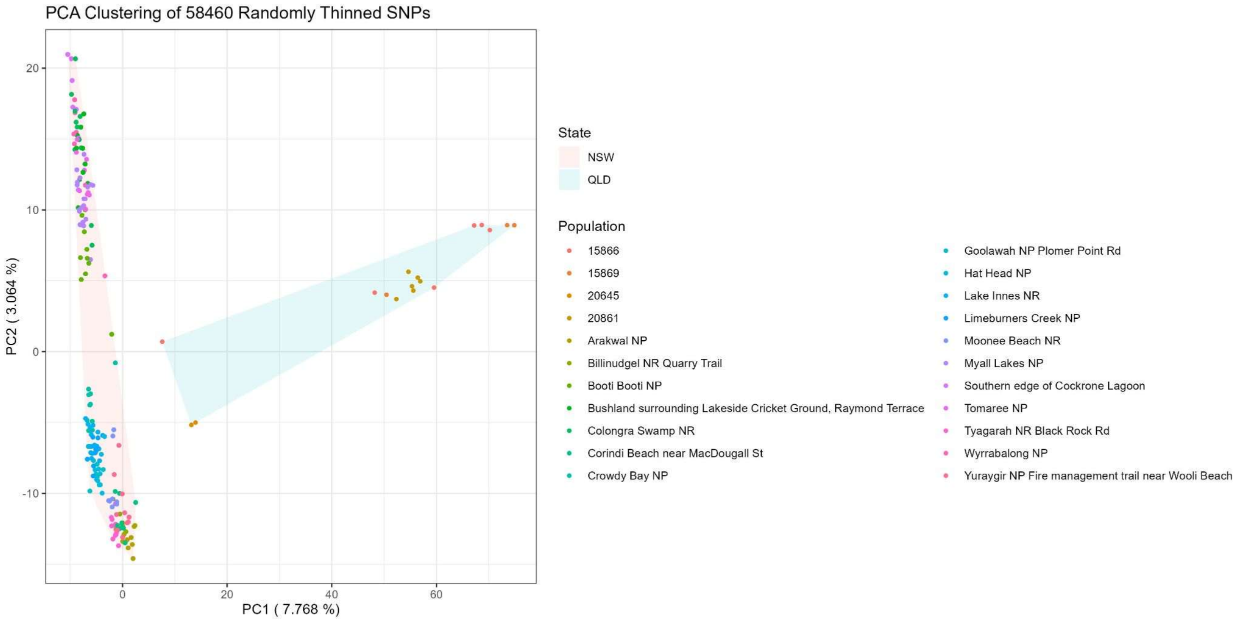


*Figure S5. Principal component analysis of the SNPs in the thinned and filtered genetic data of maternal trees of* Melaleuca quinquenervia *(NSW and QLD).*

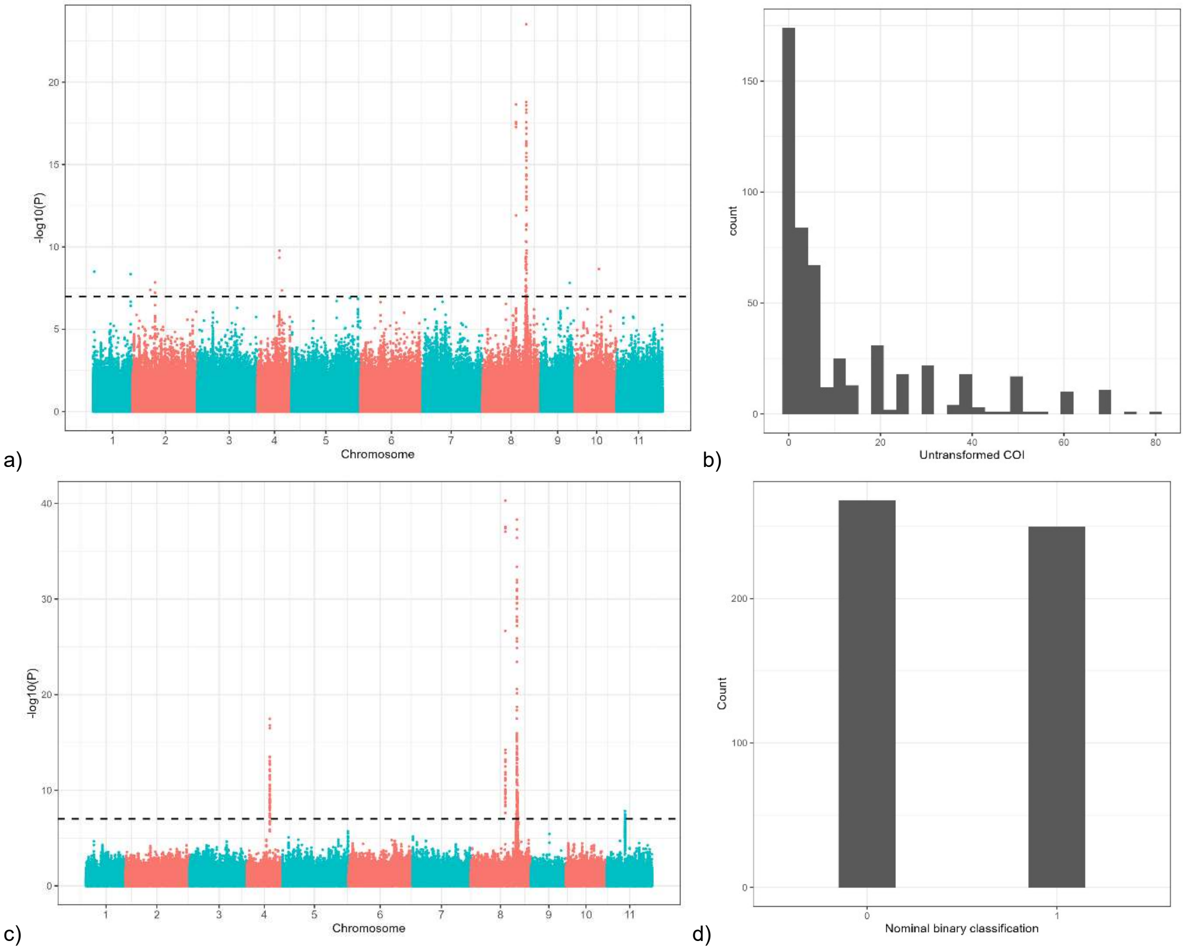


*Figure S6. Genome Wide Association Analysis of the seedling genetic dataset using different representations of phenotypes. a) Manhattan plot for seedling genetic variants and the untransformed coefficient of infection (COI) measurements. b) Frequency distribution of the untransformed COI values. c) Manhattan plot for seedling genetic variants and a binary transformation of the myrtle rust scoring scheme from the protocol described previously by Sandhu and Park (2013) - where scores between 0 and 2, inclusive, were designated a '0' and scores greater than 2 were designated a '1'. d) Frequency distribution of the binary COI values.*

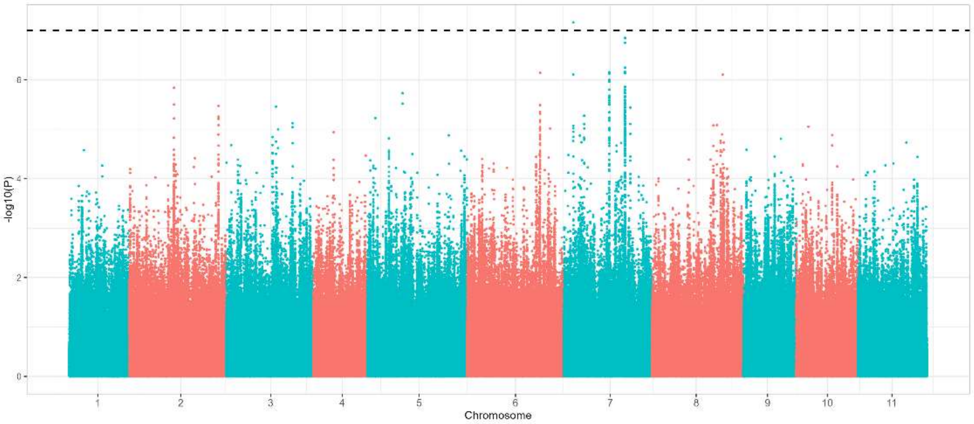


*a)*

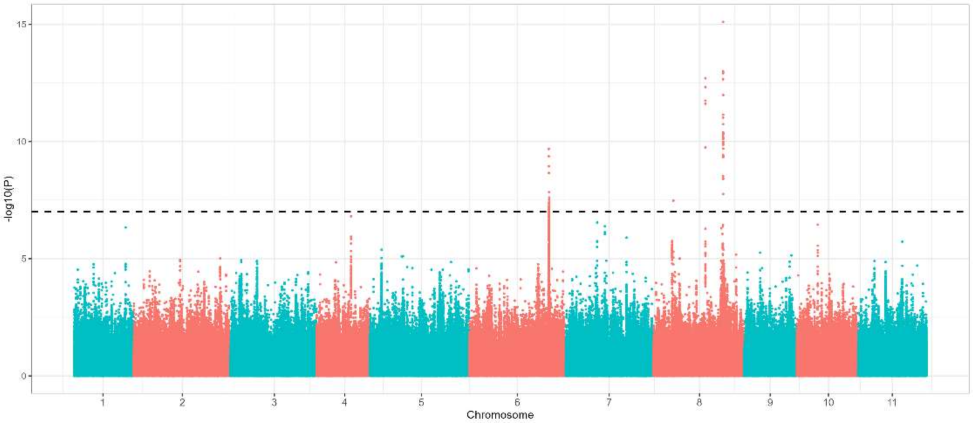


*b)*

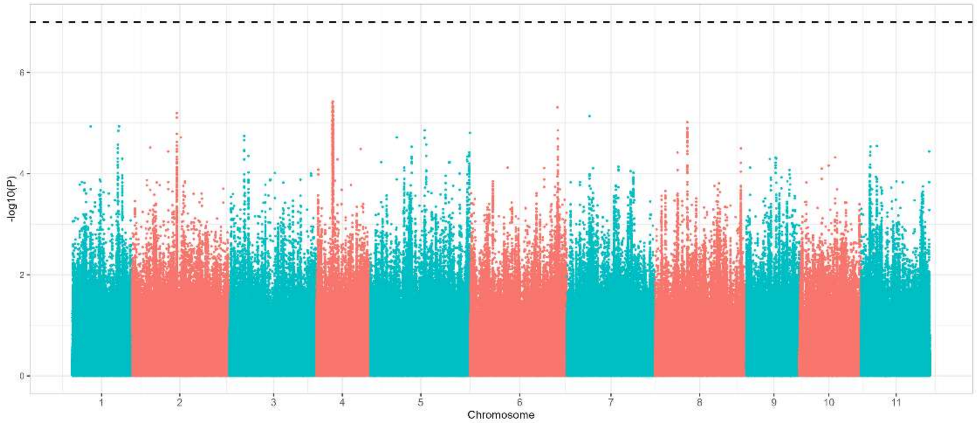


*c)*

*Figure S7. Genome Wide Association Analysis of the seedling genetic dataset for different infection and immunity phenotypes. Manhattan plot of association to immune types scored during the artificial inoculation. The horizontal dotted line marks the threshold of 1 x $10^{-7}$. a) Necrotic response, b) Flecking response, c) Chlorosis response*

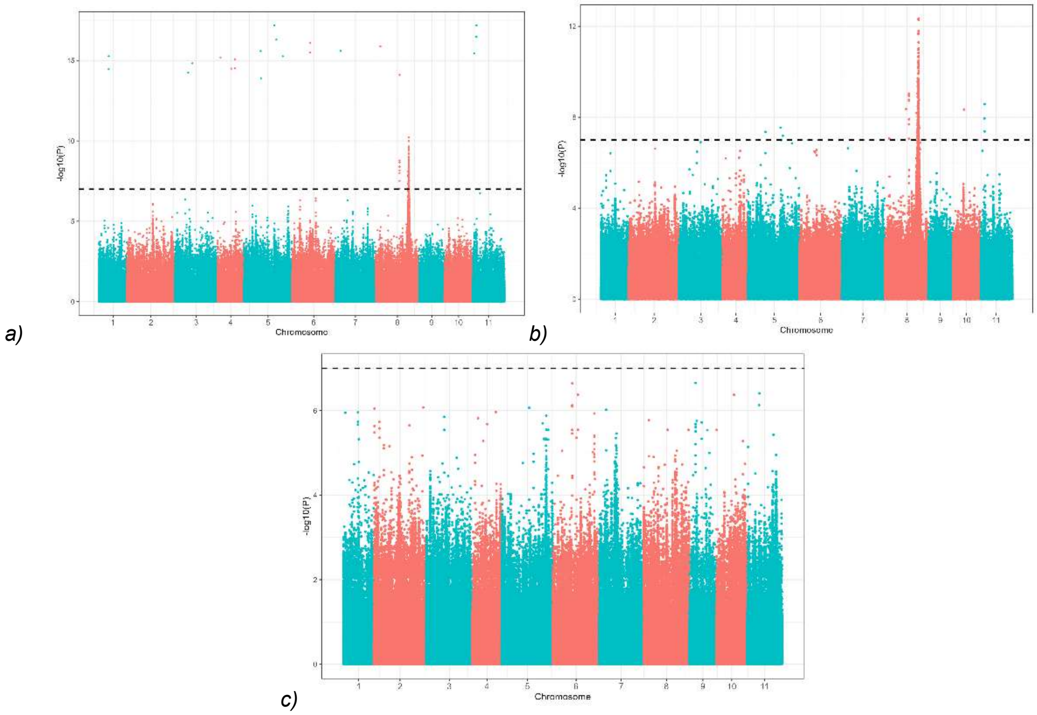


*Figure S8. Genome Wide Association Analysis of the maternal genetic dataset. Data points have been thinned to 1 in 5 SNPs for computational efficiency during plotting, with the exception of SNPs with a p-value lower than 0.01. The horizontal dotted line marks the threshold of 1 x 10-7. a) Manhattan plot between maternal genetic variants and best linear unbiased predictions of breeding values for coefficient of infection. b) Manhattan plot of association to square-root transformed mean seedling COI by family. c) Manhattan plot between maternal genetic variants and best linear unbiased predictions of breeding values for relative growth rate.*

*Table S2. Top twenty associated SNPs from the genome wide association analysis, and their associated SNPeff analysis results sorted by putative impact. Twenty-five missense variations present in the top 500 associated SNPs have also been included. Despite different gene IDs (*ID from Chen et al. *2023) on the annotated datasets, they are identical and both have been listed for ease of reference to readers. Where multiple effects were reported for one SNP, the first occurrence is reported below.*

| Chromosome 8 Position | Allele identity | Amino acid shifts | *β* | Standard error | Bonferroni adjusted p-value Wald test | Variant function (position) | Genome Annotated Gene ID |
|---|---|---|---|---|---|---|---|
| **Top 20 GWAS SNPs associated with COI using the seedling dataset** | | | | | | | |
| 15434865 | G/A | NA | -0.91907 | 0.078693 | 5.75E-28 | Intergenic region | NBARC ID: g9233-g10387 NLR ID: g4997-g6020 |
| 19357911 | G/A | NA | 0.681659 | 0.059739 | 6.69E-27 | Downstream gene variant | NBARC ID: g97 NLR ID: g9180 |
| 19336605 | G/A | Leucine to Leucine | -0.88346 | 0.078219 | 1.91E-26 | Low synonymous variant | NBARC ID: g93 NLR ID: g9176 |
| 15434549 | A/T | NA | -0.89694 | 0.079642 | 2.56E-26 | Intergenic region | NBARC ID: g9233-g10387 NLR ID: g4997-g6020 |
| 15434318 | A/G | NA | -0.89187 | 0.079631 | 4.48E-26 | Intergenic region | NBARC ID: g9233-g10387 NLRID: g4997-g6020 |
| 15435007 | A/G | NA | -0.88795 | 0.07951 | 5.98E-26 | Intergenic region | NBARC ID: g9233-g10387 NLR ID: g4997-g6020 |
| 19338387 | G/A | NA | 0.769115 | 0.069777 | 2.19E-25 | Downstream gene variant | NBARC ID: g93 NLR ID: g9176 |
| 19339421 | C/G | NA | -0.88373 | 0.080689 | 4.09E-25 | Downstream gene variant | NBARC ID: g93 NLR ID: g9176 |
| 19349317 | C/T | NA | 0.635275 | 0.059782 | 7.16E-24 | Upstream gene variant | NBARC ID: g97 NLR ID: g9180 |
| 19358240 | A/G | NA | 0.648683 | 0.061797 | 2.20E-23 | Downstream gene variant | NBARC ID: g97 NLR ID: g9180 |
| 19349473 | G/A | NA | 0.627811 | 0.059854 | 2.36E-23 | Upstream gene variant | NBARC ID: g97 NLR ID: g9180 |
| 19348127 | G/A | NA | 0.626318 | 0.059865 | 2.98E-23 | Upstream gene variant | NBARC ID: g97 NLR ID: g9180 |

| | | | | | | | |
|---|---|---|---|---|---|---|---|
| 19349286 | C/T | NA | 0.628707 | 0.060259 | 3.81E-23 | Upstream gene variant | NBARC ID: g97 NLR ID: g9180 |
| 19347992 | G/A | NA | 0.622495 | 0.059844 | 5.00E-23 | Upstream gene variant | NBARC ID: g97 NLR ID: g9180 |
| 19348984 | A/C | NA | 0.627218 | 0.06064 | 8.26E-23 | Upstream gene variant | NBARC ID: g97 NLR ID: g9180 |
| 19349526 | G/A | NA | 0.62188 | 0.060124 | 8.27E-23 | Upstream gene variant | NBARC ID: g97 NLR ID: g9180 |
| 19347539 | A/T | NA | 0.601847 | 0.058362 | 1.08E-22 | Upstream gene variant | NBARC ID: g97 NLR ID: g9180 |
| 19339802 | G/A | NA | 0.746169 | 0.074458 | 1.27E-21 | Downstream gene variant | NBARC ID: g93 NLR ID: g9176 |
| 19348203 | A/G | NA | 0.594974 | 0.059405 | 1.33E-21 | Upstream gene variant | NBARC ID: g97 NLR ID: g9180 |
| 19349112 | C/T | NA | 0.614234 | 0.061341 | 1.36E-21 | Upstream gene variant | NBARC ID: g97 NLR ID: g9180 |
| **Miss-sense variants detected in the top 500 associated SNPs** | | | | | | | |
| 19223991 | A/G | Aspartic acid to Glycine | 0.614234 | 0.061341 | 1.36E-21 | Missense-variant | NBARC ID: g82 NLR ID: g9164 |
| 19224489 | A/G | Lysine to Arginine | 0.404499 | 0.07808 | 3.24E-07 | Missense-variant | NBARC ID: g82 NLR ID: g9164 |
| 19224521 | A/G | Asparagine to Aspartic acid | 0.336765 | 0.074688 | 8.16E-06 | Missense-variant | NBARC ID: g82 NLR ID: g9164 |
| 19224562 | A/G | Isoleucine to Methionine | 0.439879 | 0.075654 | 1.10E-08 | Missense-variant | NBARC ID: g82 NLR ID: g9164 |
| 19224573 | A/G | Aspartic acid to Glycine | 0.499502 | 0.079884 | 8.80E-10 | Missense-variant | NBARC ID: g82 NLR ID: g9164 |
| 19224744 | G/A | Serine to Asparagine | 0.489552 | 0.080006 | 1.93E-09 | Missense-variant | NBARC ID: g98 NLR ID: g9164 |
| 19224780 | C/T | Threonine to Methionine | 0.421647 | 0.079431 | 1.68E-07 | Missense-variant | NBARC ID: g98 NLR ID: |

| | | | | | | | |
|---|---|---|---|---|---|---|---|
| | | | | | | | g9164 |
| 19335439 | A/C | Tyrosine to Serine | 0.430901 | 0.079529 | 9.47E-08 | Missense-variant | NBARC ID: g93 NLR ID: g9176 |
| 19335449 | T/A | Aspartic acid to Glutamic acid | 0.656111 | 0.074698 | 2.68E-17 | Missense-variant | NBARC ID: g93 NLR ID: g9176 |
| 19336685 | C/T | Proline to Leucine | 0.656111 | 0.074698 | 2.68E-17 | Missense-variant | NBARC ID: g93 NLR ID: g9176 |
| 19336702 | G/A | Glycine to Arginine | 0.722016 | 0.07493 | 3.09E-20 | Missense-variant | NBARC ID: g93 NLR ID: g9176 |
| 19336837 | T/G | Tyrosine to Aspartic acid | 0.382113 | 0.082233 | 4.34E-06 | Missense-variant | NBARC ID: g93 NLR ID: g9176 |
| 19336921 | A/G | Isoleucine to Valine | 0.689064 | 0.073069 | 1.64E-19 | Missense-variant | NBARC ID: g93 NLR ID: g9176 |
| 19337434 | A/G | Lysine to Glutamic acid | 0.370074 | 0.086014 | 2.04E-05 | Missense-variant | NBARC ID: g93 NLR ID: g9176 |
| 19337446 | A/G | Threonine to Alanine | 0.384517 | 0.084739 | 7.17E-06 | Missense-variant | NBARC ID: g93 NLR ID: g9176 |
| 19337681 | A/G | Lysine to Arginine | 0.412216 | 0.094723 | 1.65E-05 | Missense-variant | NBARC ID: g93 NLR ID: g9176 |
| 19337758 | T/C | Phenylalanine to Leucine | 0.323259 | 0.077041 | 3.23E-05 | Missense-variant | NBARC ID: g93 NLR ID: g9176 |
| 19337785 | A/T | Methionine to Leucine | 0.717154 | 0.074891 | 5.05E-20 | Missense-variant | NBARC ID: g93 NLR ID: g9176 |
| 19337790 | A/T | Arginine to Serine | 0.714175 | 0.074925 | 7.23E-20 | Missense-variant | NBARC ID: g93 NLR ID: g9176 |
| 19337815 | A/G | Lysine to Glutamic acid | 0.404335 | 0.080926 | 8.15E-07 | Missense-variant | NBARC ID: g93 NLR ID: g9176 |
| 19511746 | A/G | Isoleucine to Valine | 0.320912 | 0.074166 | 1.83E-05 | Missense-variant | NBARC ID: g116 NLR ID: g9199 |
| 19511755 | G/A | Aspartic acid to Asparagine | -0.4222 | 0.070809 | 4.76E-09 | Missense-variant | NBARC ID: g116 NLR ID: |

| | | | | | | | |
|---|---|---|---|---|---|---|---|
| | | | | | | | g9199 |
| 19512320 | T/A | Tyrosine to Asparagine | 0.3997 | 0.09325 | 2.19E-05 | Missense-variant | NBARC ID: g116 NLR ID: g9199 |
| 19513767 | T/C | Leucine to Proline | -0.41577 | 0.071067 | 8.99E-09 | Missense-variant | NBARC ID: g116 NLR ID: g9199 |
| 19513854 | C/T | Serine to Leucine | 0.377403 | 0.084204 | 9.22E-06 | Missense-variant | NBARC ID: g116 NLR ID: g9199 |

*Table S3. Results of ABBA-BABA analyses across the whole genome. D-statistic values annotated with asterix to indicate size of the values (>0.01*, >0.1**, >0.15***). Bold p-values represent values below 0.05.* Melaleuca quinquenervia *populations from NSW were split into north and south (threshold of -29 °S latitude). Samples were chosen at random to represent these regions and to represent Queensland. Among the ABBA-BABA analyses presented below, three different topologies were inferred between populations of M. quinquenervia from the south (qsouth), a population of* M. quinquenervia *from further north (qnorth), a representative of different broadleaf Melaleuca lineage (*M. viridiflora, M. argentea, M. cajuputi or M. leacadendra*: collectively vacl), and outgroup* M. nodosa *(out). These are illustrated below as scenarios. Each ABBA-BABA analysis tests for an excess of derived allele sharing between the designated P2 and P3 populations.*

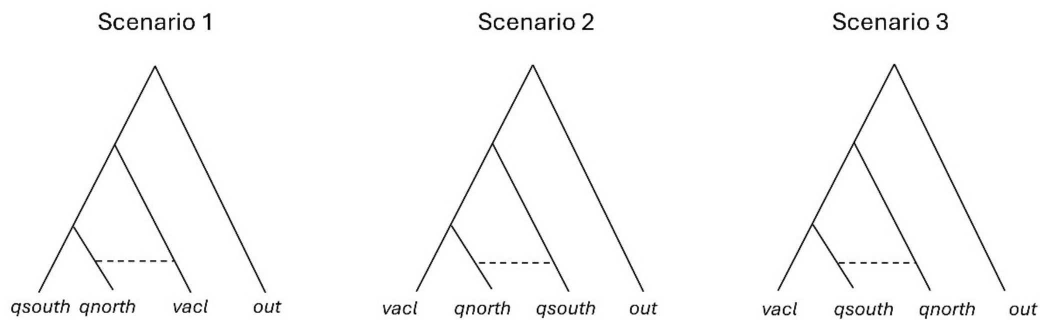


| P1 | P2 | P3 | Scenario | D-statistic | Z-score | p-value | f4-ratio | BBAA | ABBA | BABA |
|---|---|---|---|---|---|---|---|---|---|---|
| M. quin. SNSW | M. quin. QLD | M. viridiflora | 1 | 0.189132*** | 12.9058 | **2.30E-16** | 0.077686 | 144450 | 107724 | 73456.8 |
| M. quin. NNSW | M. quin. QLD | M. viridiflora | 1 | 0.170297*** | 12.6843 | **2.30E-16** | 0.071031 | 146638 | 106886 | 75779 |
| M. quin. SNSW | M. quin. NNSW | M. viridiflora | 1 | 0.024305* | 5.49394 | **3.93E-08** | 0.00717 | 200283 | 66621 | 63459.4 |
| M. quin. SNSW | M. quin. QLD | M. argentea | 1 | 0.157549*** | 10.9801 | **2.30E-16** | 0.037681 | 183031 | 77261.5 | 56230 |
| M. quin. NNSW | M. quin. QLD | M. argentea | 1 | 0.138482** | 9.85731 | **2.30E-16** | 0.033619 | 185558 | 76817.7 | 58129.9 |
| M. quin. SNSW | M. quin. NNSW | M. argentea | 1 | 0.02357* | 13.0945 | **2.30E-16** | 0.0042 | 243002 | 50889.2 | 48545.6 |
| M. cajuputi | M. quin. QLD | M. quin. SNSW | 2 | 0.168541*** | 14.3171 | **2.30E-16** | 0.260291 | 129757 | 117041 | 83278.6 |
| M. cajuputi | M. quin. QLD | M. quin. NNSW | 2 | 0.158782*** | 13.6674 | **2.30E-16** | 0.269104 | 128863 | 118801 | 86243.9 |
| M. quin. SNSW | M. quin. NNSW | M. cajuputi | 1 | 0.026114* | 7.93749 | **2.06E-15** | 0.010387 | 173747 | 75862.4 | 72001.1 |

| | | | | | | | | | | |
|---|---|---|---|---|---|---|---|---|---|---|
| M. leucadendra | M. quin. SNSW | M. quin. QLD | 3 | 0.001 214 | 0.1724 85 | **0.8630 56** | 0.0026 77 | 14157 2 | 11846 7 | 11818 0 |
| M. leucadendra | M. quin. NNSW | M. quin. QLD | 3 | 0.014 075* | 2.0074 1 | **0.0447 06** | 0.0313 19 | 14495 5 | 12108 7 | 11772 6 |
| M. quin. SNSW | M. quin. NNSW | M. leucadend ra | 1 | 0.018 835* | 5.1215 5 | **3.03E-07** | 0.0131 34 | 15370 9 | 10372 4 | 99888. 9 |

145

#### Seedling COI GWAS with different filters and analytical methods

Different variant filtering combinations were used for GWAS analyses to ensure results were not highly sensitive to these filters. These included varying the mapping quality filter, variant filters including site missingness, departure from Hardy-Weinberg equilibrium, and minor allele frequencies, and the model used for GWAS (linear mixed model or Bayesian sparse linear mixed model). For each analysis, the top effect scores (*p-value* < 0.01 for linear-mixed model runs and top 1% effect for Bayesian sparse linear mixed models) were plotted as a Manhattan plot. We note that some parameters resulted in changes in the density of associated SNPs. However, across many different combinations of parameter values and model inference methods, a common set of regions exhibited strong associations.

We note that the 1 technical replicate remaining in the seedling dataset was not removed for the following iterations of GWAS analyses.

*Table S4. Table of iterations of analyses conducted with different filters on the seedling dataset to determine the sensitivity of significantly associated MR resistance loci. In bold are the key changes made to the filters used in the iteration compared to the previous. Abbreviations: LMM = Linear Mixed Model; BSLMM = Bayesian Sparse Linear Mixed Model*

| Iteration | Mapping Quality Filter | Read Depth (DP) | Genotyping Quality (GQ) | Missingness | Hardy-Weinberg Equilibrium | Mean sample read depth (sample filter) | COI Phenotype input | Minor allele frequency | GWAS method | Top effect score Manhattan plots |
|---|---|---|---|---|---|---|---|---|---|---|
| 1 | None | 6 | 20 | 150 | None | 4 | Raw | 0.01 | LMM | 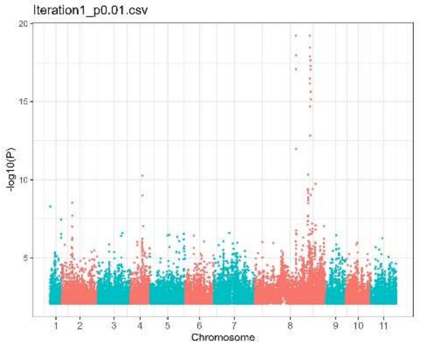  |

| | | | | | | | | | | |
|---|---|---|---|---|---|---|---|---|---|---|
| 2 | None | 6 | 20 | **200** | None | 4 | **Square root** | 0.01 | LMM | 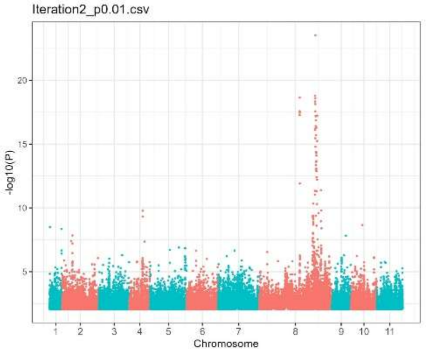 |
| 3 | None | 6 | 20 | 200 | None | 4 | **Quarter power** | 0.01 | LMM | 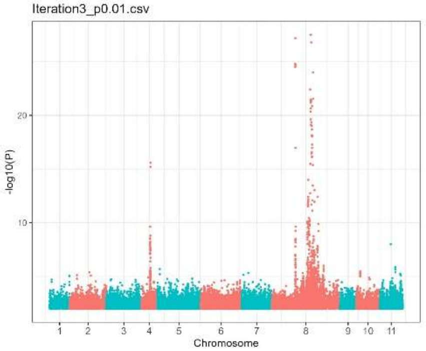 |
| 4 | None | 6 | 20 | 200 | None | 4 | Quarter power | **0.05** | LMM | 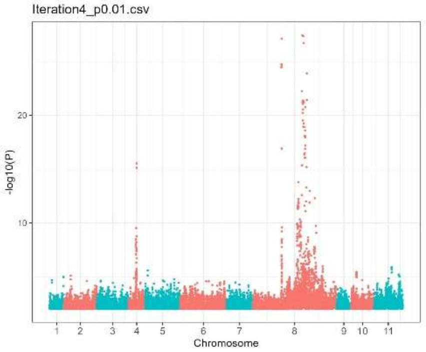 |

| | | | | | | | | | | |
|---|---|---|---|---|---|---|---|---|---|---|
| 5 | None | 6 | 20 | 200 | **1E-5** | 4 | Quarter power | 0.05 | LMM | 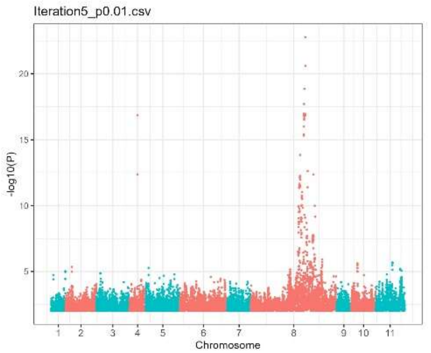 |
| 6 – chromosome 8 only | **5** | 6 | 20 | 200 | None | 4 | Quarter power | 0.05 | LMM | 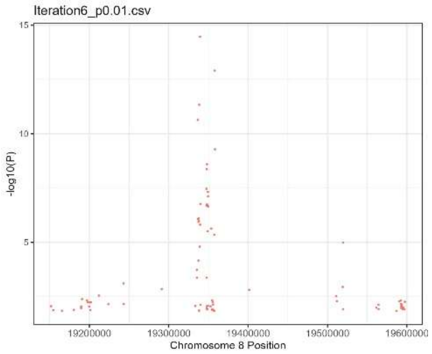 |
| 7 – chromosome 8 only | **10** | 6 | 20 | 200 | None | 4 | Quarter power | 0.01 | LMM | 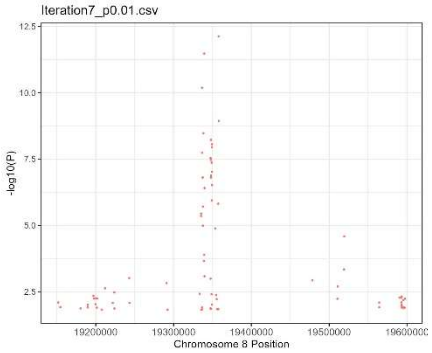 |

| | | | | | | | | | | |
|---|---|---|---|---|---|---|---|---|---|---|
| 8 – chromo some 8 only | **15** | 6 | 20 | 200 | None | 4 | Quarter power | 0.01 | LMM | 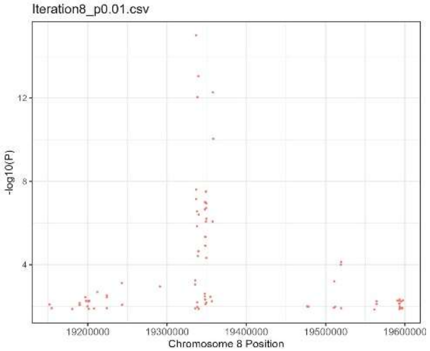 |
| 9 – chromo some 8 only | **20** | 6 | 20 | 200 | None | 4 | Quarter power | 0.01 | LMM | 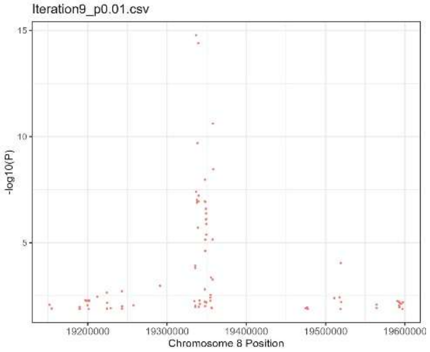 |
| 10 – chromo some 8 only | **30** | 6 | 20 | 200 | None | 4 | Quarter power | 0.01 | LMM | 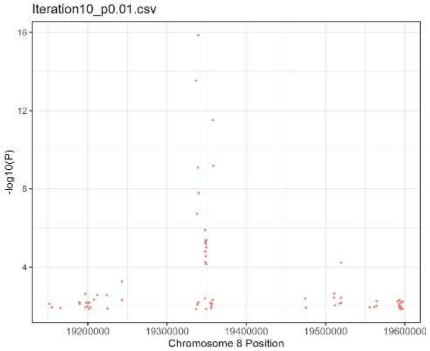 |

| | | | | | | | | | | |
|---|---|---|---|---|---|---|---|---|---|---|
| 11 (iter_4/ gwas) | 15 | 6 | 20 | 200 | **1E-5** | 4 | Quarter power | 0.05 | LMM | 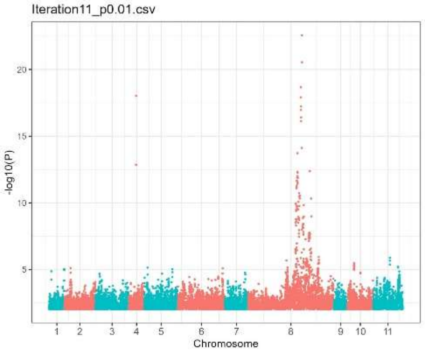 |
| 12 – FINAL | 15 | 6 | 20 | 200 | **None** | 4 | Quarter power | 0.05 | LMM | 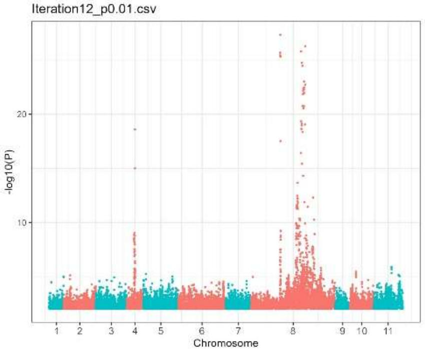 |
| 13 | 15 | 6 | 20 | 200 | None | 4 | **Binary phenotype where MR Immune Scoring <=2C vs 2C>** | 0.05 | LMM | 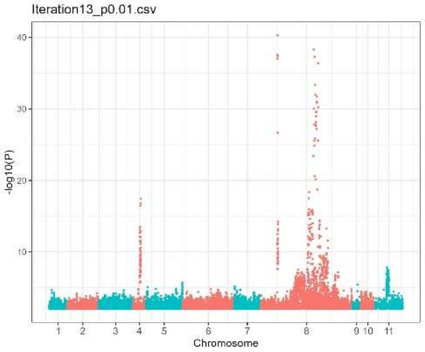 |

| | | | | | | | | | | |
|---|---|---|---|---|---|---|---|---|---|---|
| 14 | 15 | 6 | 20 | 200 | None | 4 | **Binary phenotype where MR Immune Scoring 0 vs 1+** | 0.05 | LMM | 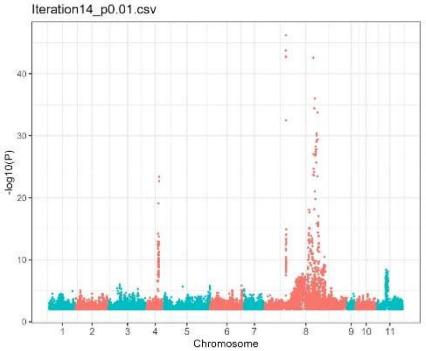 |
| 15 | 15 | 6 | 20 | 200 | None | 4 | **Binary phenotype - Chlorotic immune response presence** | 0.05 | LMM | 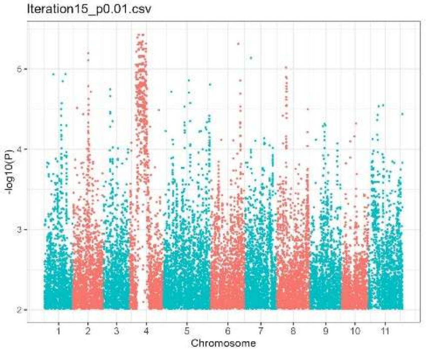 |
| 16 | 15 | 6 | 20 | 200 | None | 4 | **Binary phenotype - Necrosis immune response presence** | 0.05 | LMM | 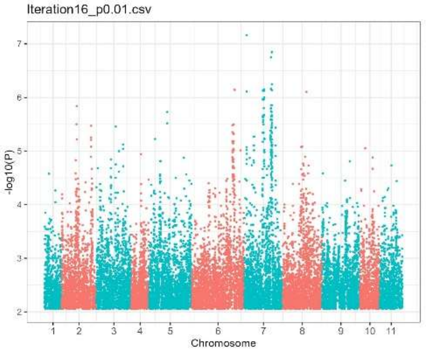 |

| | | | | | | | | | | |
|---|---|---|---|---|---|---|---|---|---|---|
| 17 | 15 | 6 | 20 | 200 | None | 4 | **Binary phenotype - Flecking immune response presence** | 0.05 | LMM | 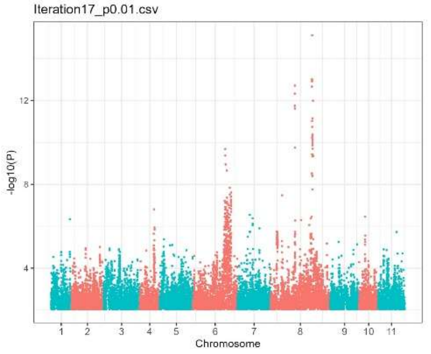 |
| 18 | 15 | 6 | 20 | 200 | 1E-5 | 4 | Quarter power | 0.05 | **BSLMM** | 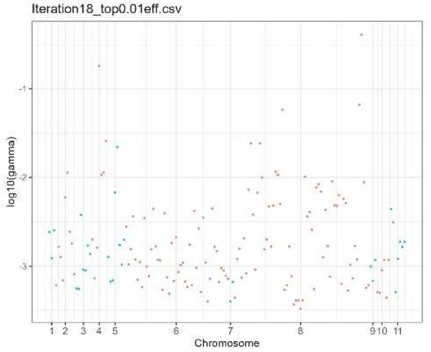 |
| 19 | 15 | 6 | 20 | 200 | 1E-5 | 4 | Quarter power | 0.05 | **BSLMM, increase steps 10x (1E7) and burn-in 20% steps (2E6)** | 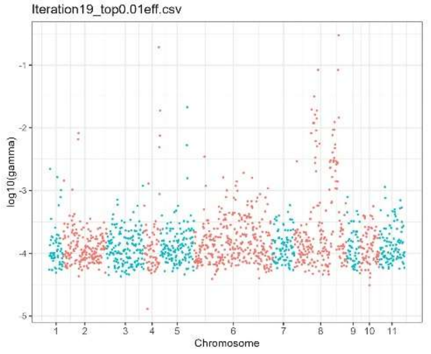 |

#### Gene annotations in a key region of Chromosome 8

To better understand variation in a region on Chromosome 8 that featured a large number of SNPs associated with resistance, we examined NLR genes annotated on Haplotypes A and B of the phased reference genome. Genes detected in this region of Haplotype A (between g93, ~19.33 million bp, and g98, ~19.37 million bp) were BLAST-ed (Altschul et al., 1990; Zhang et al., 2000, Camacho et al. 2009) to Haplotype B and filtered by bitscore (> 3,000). These produced an initial set of regions with high levels of identity between the haplotypes. These regions were checked using Anchorwave gapped alignments (Song et al. 2022) of Chromosome 8 to confirm broad synteny of regions and to confirm putative deletions or insertions between Haplotypes A and B. The regions that were putatively matched by BLAST were then manually matched to an overlapping NBARC gene ID in the annotation on the genome (Chen et al. 2023). In addition, for each gene of interest on Haplotype A, a protein-to-protein BLAST (BLASTp) was performed with the Haplotype A gene as the query, and the NBARC gene annotations of Haplotype B as the database. BLASTp results were filtered for the top 3 strongest matching proteins on Chromosome 8. These provided three indications for correspondence between sequences on Haplotypes A and B. Results on Haplotype B were filtered to Chromosome 8 from position 19,989,090 to position 20,223,658, informed by the Anchorwave alignment. Gene MQUIG19794 and MQUIG19795 (Chen et al. 2023) in this region of Haplotype A did not match any genes on Haplotype B. When the latter was BLAST to the NCBI database, it returned strong hits to reverse transcriptase (NCBI name RVT-2, NCBI accession pfam07727, query interval 1-109, E-value = 8.87E-25; NCBI name RNASE_HI_RT_Ty1, NCBI accession cd09272, query interval 195-331, E-value = 1.50E-72). The genes of interest and their putative matches were then plotted using gggenomes (Hackl et al. 2024) along with the transposons found in the regions.

#### Population-wide haplotype variation in Gene 97

Some of the SNPs that we observed to be most strongly associated with myrtle rust resistance occurred in the region of Gene 97 (Chromosome 8, Haplotype A, position 19,350,017 to 19,354,857). We extracted the alignment for this region from the phased VCF, and considered allelic variations in the inferred protein sequences. In summary, there were 464 unique sequences across the haplotypes of our seedling dataset (Fig. 6a). Within some variations of the amino acid sequences, stop codons were encoded prematurely at positions 931, 985 and 1329. Broadly we observed relatively little clustering of resistance phenotypes with the inferred amino acid sequence of haplotypes (Fig. 6b). We note that phenotype-genotype links could be obscured by dominance in this analysis, which assigns one phenotype to each of the inferred haplotypes from an individual. However, these inferred sequences might be useful later for understanding links between sequence variation and phenotypes.

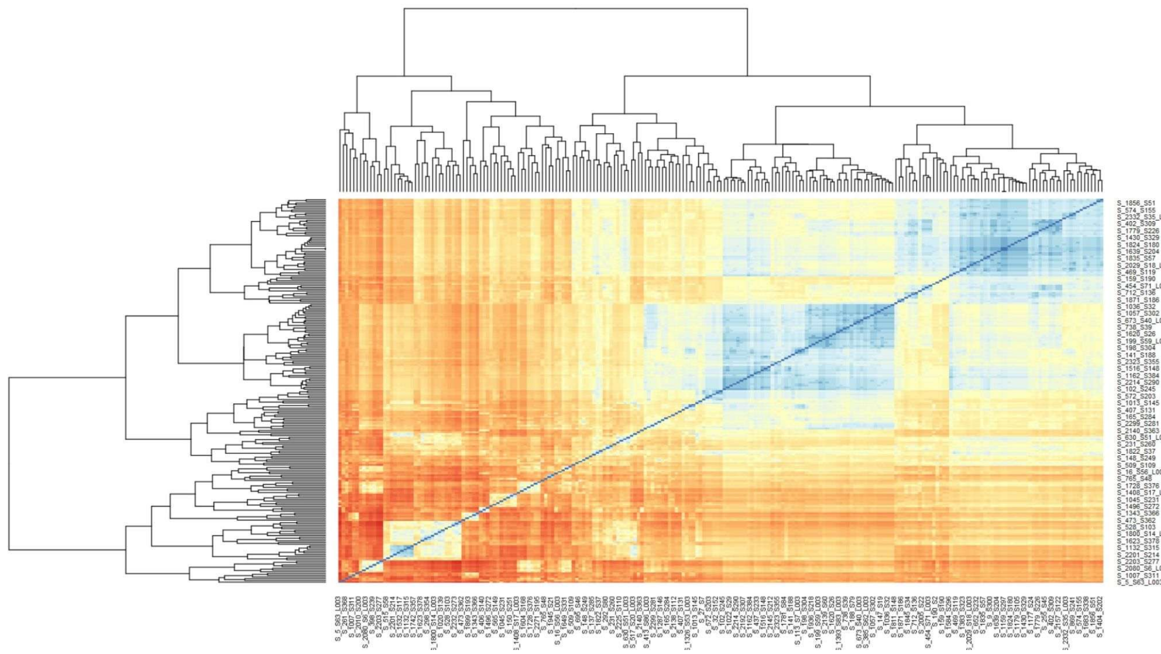

a)

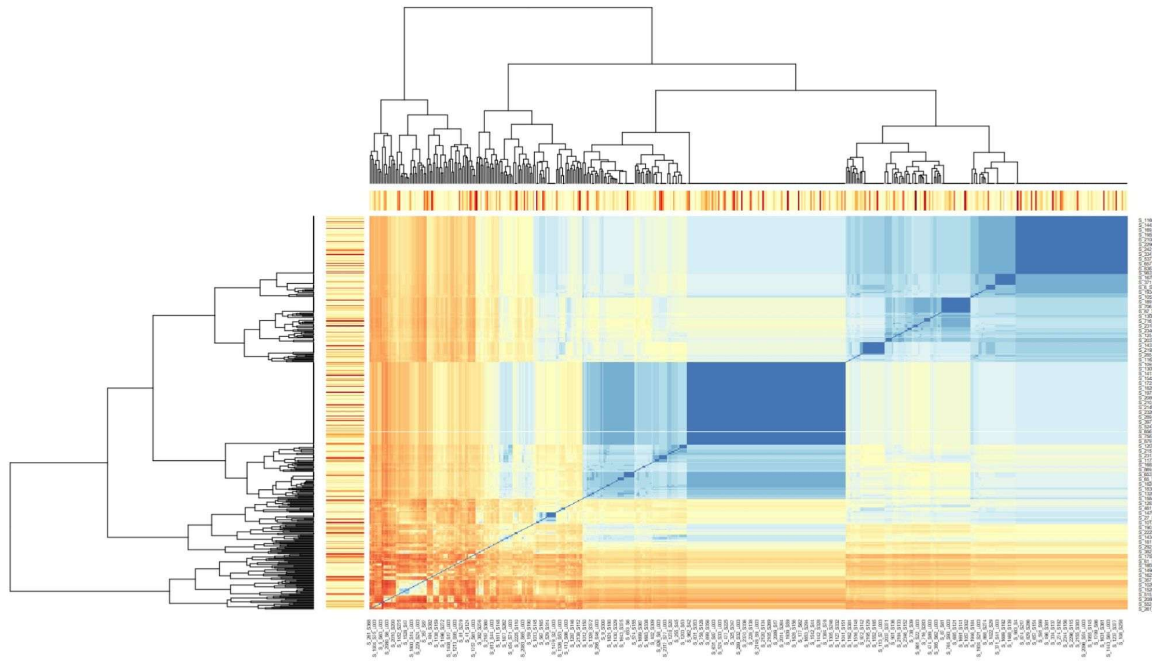

b)

*Figure S9. Heatmap of gene 97's distance matrix when comparing two individual's allelic series. a) Only representing individuals that are unique. b) Individuals with identical sequences are included. Coloured bars on the top and left-side of the heatmap represent the individual's coefficient of infection, with red representing a low resistance individual (high COI) and yellow representing high resistance (low COI).*

#### Genomic prediction

<u>Genomic Prediction Iterations</u>

The goal of our study was to create a targeted genotyping protocol that would enable us to characterise susceptibility to myrtle rust in plants in the most cost-efficient manner. To find an approach that was effective, we fit multiple iterations of genomic prediction models while shifting variables including: the number of SNPs used in the model, the filters applied to the SNPs, use of different phenotypes, and thinning of SNPs. We also used different approaches to test the robustness of cross-validation (Table S5-6), including the use of outlier populations and families to check the model's accuracy across the geographical range examined (Table S9). We outline the results of a few iterations here to highlight some useful principles.

The final iteration (subsequently also referred to as Iteration 48) used a mix of 999 MR significant SNPs thinned by 1 SNP per 1 kbp window, and 50 most strongly associated epistatic SNPs, totalling to 1,049 SNPs. The iteration had a reliability of 0.683, and an $R^2$ of 0.683 between testing individuals (R=0.83 2 D.P.). Further resampling of randomised populations and maternal lines used for training/testing are reported below.

*Table S5. Randomly selecting 35 maternal lines used for testing. Maternal lines were sourced from the seedling's maternal lines locations. For each iteration, three maternal lines were excluded from the training data, these varied in sizes thus varying the training size.*

| Training number | $R^2$ testing individuals |
|---|---|
| 417 | 0.632816 |
| 422 | 0.621449 |
| 423 | 0.721841 |
| 399 | 0.679173 |
| 412 | 0.663161 |

*Table S6. Maternal lines were grouped by population in accordance to location sampled. Randomising the populations used for testing revealed robust $R^2$ accuracy within the predicted and ground-truth resistance against myrtle rust. For each iteration, three populations were excluded from the training data, these populations varied in sizes thus varying the training size.*

| Testing populations | Training Number | $R^2$ testing individuals |
|---|---|---|
| p72bbeb40, popnInfo-w4ba03a, popnInfo-m1lg07j | 442 | 0.692307 |
| pbb515550, p9c3c9590, pa9710de0 | 449 | 0.687889 |
| popnInfo-m1lg07j, p72bbeb40, p328a8440 | 421 | 0.679417 |
| popnInfo-w4ba03a, pd135bc30, p831064a0 | 452 | 0.684642 |
| p328a8440, p9c3c9590, p1bff81d0 | 435 | 0.668776 |

Varying the number of strongly associated SNPs

The costs to develop and use a targeted genotyping protocol typically increases with the number of markers involved. As such, we wanted to determine the smallest number of SNPs we could use in a model to successfully quantify susceptibility in a satisfactory manner. SNP numbers fewer than 500 were insufficient, with poor $R^2$ values when cross-validated against ground-truth COI values (Iteration 1-4). Likewise, we observed decreased model quality with large numbers of SNPs (Iteration 31, 17). This is further explored in the section detailing iterations using thinned SNPs.

The impacts of different data filters were also explored (corresponding to GWAS results in Table S4). For example, removing SNPs that exhibited excessive departure from Hardy Weinberg allele proportions sometimes reduced the predictive power of models (Iterations 17 and 31, Table S4). Iterations 27-30 explored how thinning of the strongly associated SNPs impacted the model. Thinning was conducted by removing associated SNPs at differing window sizes. Interestingly, the thinning of these SNPs resulted in improved model predictions (e.g. comparing Iteration 17 to Iteration 28, based on the results of the individuals held out for cross-validation). This model improvement was observed until SNPs were thinned at a 2 kbp window size (Iteration 30), where a slight decrease in model quality was observed.

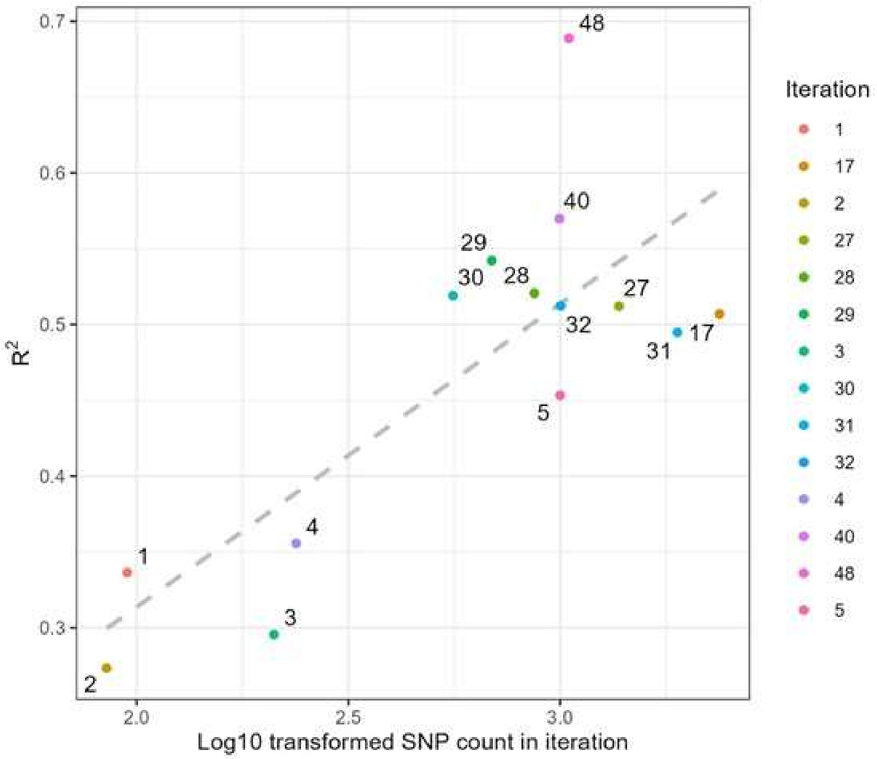


*Figure S10. A select few iterations to represent the relationship between the number of SNPs and the $R^2$ value between predicted COI and the ground-truth COI for individuals that were not used to train the genomic prediction model. The dashed line represents a linear model between the number of SNPs and their corresponding $R^2$ value.*

*Table S7. The iteration number, the number of SNPs used in the genomic prediction model for the iteration, a description of the SNPs used, the reliability of the predictions made and the $R^2$ value of all individuals (training and cross-validation individuals combined). Here, the iterations focus on changing the number of SNPs used and varying the filter of the genetic data.*

| Iteration | SNP count | Description | Mean reliability | $R^2$ |
|---|---|---|---|---|
| 1 | 95 | Using SNPs from the GWAS. Genetic data is filtered as described in the manuscript, with a mapping quality filter of 15. | 0.034132 | 0.3363477 |
| 2 | 85 | Using the top 85 most strongly associated SNPs. Genetic data is filtered as described in the manuscript, with a mapping quality filter of 15. | 0.050525 | 0.2732518 |
| 3 | 211 | Using SNPs with a p-value of less than 1E-4 from the GWAS. Genetic data is filtered as described in the manuscript, with a mapping quality filter of 15 and a Hardy-Weinberg filter of 1E-4. | -0.14425 | 0.2954709 |
| 4 | 238 | Using SNPs with a p-value of less than 1E-4 from the GWAS. Genetic data is filtered as described in the manuscript, with a mapping quality filter of 15. | -0.10139 | 0.3556766 |
| 5 | 1000 | Using the top 1,000 SNPs from the GWAS. Genetic data is filtered as described in the manuscript, with a mapping quality filter of 15 and a Hardy-Weinberg filter of 1E-4. | 0.023669 | 0.4532515 |
| 17 | 2377 | Using SNPs with a p-value of less than 1E-3 from the GWAS. Genetic data is filtered as described in the manuscript, with a mapping quality filter of 15. | 0.372008 | 0.5069734 |
| 27 | 1377 | Using the SNPs in Iteration 17, and thinning by 1 SNP per 100bp | 0.502004 | 0.5121157 |
| 28 | 869 | Using the SNPs in Iteration 17 and thinning by 1 SNP per 500bp | 0.611174 | 0.5206568 |
| 29 | 689 | Using the SNPs in Iteration 17 and thinning by 1 SNP per 1,000bp | 0.634427 | 0.5421888 |
| 30 | 559 | Using the SNPs in Iteration 17 and thinning by 1 SNP per 2,000bp | 0.66333 | 0.5191254 |
| 31 | 1890 | Using the SNPs in Iteration 17 and placing a Hardy-Weinberg filter of 1E-4 | 0.380826 | 0.4948388 |
| 32 | 1003 | Using the SNPs in Iteration 17 and placing a Hardy-Weinberg filter of 1E-4 and thinning by 1 SNP per 200bp | 0.500022 | 0.5122733 |
| 40 | 996 | Myrtle rust associated SNPs from the original GWAS, thinned at 1 SNP per 100 bp. | 0.6298315 | 0.5697814 |

#### Genomic prediction of seedling phenotype using associated SNPs sourced from maternal lines

We next performed a series of analyses to examine whether we could use positions discovered in the maternal GWAS analysis to retrieve seedling genotypes for the genomic prediction model for prediction. If possible, this will potentially avoid the need to phenotype each seedling for future GWAS. Our first set of analyses tested whether the associated SNPs from the maternal line GWAS could be used to generate a genomically estimated breeding value of seedlings (Iteration 15, 19-21). We then reversed this analysis and determined whether the seedling's associated SNPs could be used to predict the calculated breeding values in the maternal dataset (Iteration 26).

*Table S8. The iteration number, the number of SNPs used in the genomic prediction model for the iteration, a description of the SNPs used, the mean reliability of the predictions made and the $R^2$ value of all individuals (training and cross-validation individuals combined). Here, the iterations focus on the markers discovered in the GWAS of the maternal genetic dataset and its application on predicting seedling phenotypes.*

| Iteration | SNP count | Description | Mean reliability | $R^2$ |
|---|---|---|---|---|
| 15 | 7942 | Predicting seedling phenotypes. Using SNPs with a p-value less than 1E-4 (209 SNPs) from the maternal GWAS and using thinned SNPs across the seedling genome for kinship association. | 0.37321084 | 0.1014275 |
| 19 | 524 | Repeating iteration 15. A Hardy-Weinberg threshold filter of 1 E-5 was applied on the seedling dataset. | 0.04347915 | 0.2955050 |
| 20 | 675 | Predicting seedling phenotypes. Using the top 2,000 associated SNPs of the GWAS from the maternal genetic dataset and their calculated breeding values. Only 675 SNPs were overlapping the seedling genetic dataset. This dataset was used to predict seedling resistance. | 0.19116975 | 0.2873707 |
| 21 | 9657 | Predicting seedling phenotypes. Using SNPs with a p-value less than 1 E-3 (1,933 SNPs) from the maternal GWAS that was run on the maternal genetic dataset with an additional 1 E-5 filter of Hardy-Weinberg. Thinned SNPs across the seedling genome were also added for kinship association. | 0.31602668 | 0.1606245 |

#### Training dataset validation

Genomic prediction models can be impacted by the training dataset used. This can include the effects of the size of the dataset, as well as the presence of population structure (covered in McGaugh et al., 2021; Tan et al., 2017). To test that our models were robust to these effects, we varied the volume and source of the testing and training data that were used. First, we varied the ratio of samples included in the training and testing datasets. As this ratio increased, we observed a slight but near linear decrease in the fit between predicted and observed COI ($R^2$), though overall, the accuracy of the model was sufficient for our purposes.

Second, genomic prediction models are often constrained by population structure of the species (McGaugh et al. 2021). To test the effects of this, we strategically held out populations for testing that occurred at the ends of the examined spatial range and used the remaining individuals as the training dataset. The high $R^2$ value between the ground truth and predicted COI values of these populations in tests suggested that population structure did not have a strong effect on the performance of the model, and performance was sufficient at the extremities of the examined range.

*Table S9. The iteration number, the number of SNPs used in the genomic prediction model for the iteration, a description of the SNPs used, the mean reliability of the predictions made and the $R^2$ value of all individuals (training and cross-validation individuals combined)/only cross-validation individuals. Here, the iterations focused on varying the source and volume of the testing/training datasets.*

| Iteration | SNP count | Description | Mean reliability | $R^2$ |
|---|---|---|---|---|
| 22 | 2377 | Varying the volume of testing and training individuals.<br><br>Testing:Training individuals<br>1. 100:420<br>2. 200:320<br>3. 300:220 | 1. 0.6370096<br>2. 0.7374813<br>3. 0.7196880 | 1. 0.8598871<br>2. 0.8213648<br>3. 0.7398323 |
| 25 | 2377 | Predicting Iteration 17 onto outlier populations of the north-most and south-most populations (152 testing, 368 training) | 0.627657 | 0.826054 |
| 49 | 1049 | Predicting Iteration 48 (final model selected) onto outlier populations of the north-most and south-most populations.<br>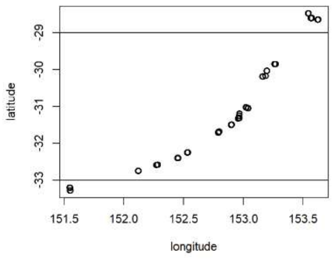 | | |

| | | | | |
|---|---|---|---|---|
| | | Using the north-most populations as training (n=405) and remaining as testing (n=87) | 0.6863841 | 0.8335309<br><br>Testing individuals only: 0.6714687 |
| | | Using the south-most populations as training (n=382) and remaining as testing (n=110) | 0.6728118 | 0.8543414<br><br>Testing individuals only: 0.7257789 |

<u>Thinned 'neutral' SNPs</u>

This set of analyses focus on using 'neutral' SNPs from the genome that are thinned using varying window sizes and examining their impact when in conjunction with associated SNPs. Iterations 12 and 33 were conducted using thinned 'neutral' and associated SNPs. The addition of the thinned 'neutral' SNPs to previous model iterations unexpectedly showed a decrease in respective model $R^2$ values (e.g., Iteration 4 & 12, Iteration 29 & 33). Which potentially may be due to the thinned SNPs contributing excess noise to the model rather than improving kinship estimations as we expected. We then examined the use of solely thinned 'neutral' SNPs in Iterations 10, 13, and 16. This resulted in a lack of increase of model predictive power despite an increasing number of random thinned SNPs, underscoring validating the necessity of using associated marker SNPs.

*Table S10. The iteration number, the number of SNPs used in the genomic prediction model for the iteration, a description of the SNPs used, the mean reliability of the predictions made and the $R^2$ value of all individuals (training and cross-validation individuals combined). Here, the iterations focused on the inclusion of neutral SNPs from thinning the genome.*

| Iteration | SNP count | Description | Mean reliability | $R^2$ |
|---|---|---|---|---|
| 4 (comparison) | 238 | Using SNPs with a p-value of less than 1E-4 from the GWAS. Genetic data is filtered as described in the manuscript, with a mapping quality filter of 15. | -0.10139 | 0.3556766 |
| 10 | 11504 | Thinning the genetic dataset by 1 SNP per 10,000 bp window. A minor allele frequency of 0.1 and a mapping quality filter of 15 were also used. | 0.486322 | 0.220512 |
| 12 | 7943 | Thinning the genetic dataset by 1 SNP per 3,000 bp window. A minor allele frequency of 0.1 and a mapping quality filter of 15 were also used. The resistance associated SNPs from iteration 4 (238 SNPs) were also used. | 0.4028624 | 0.25801528 |
| 24 | 7733 | Repeating iteration 12 with only its thinned SNPs | 0.0974314 | -0.01861275 |
| 13 | 4210 | Thinning the genetic dataset by 1 SNP per 6,000 bp window. A minor allele frequency of 0.1 and a mapping quality filter of 15 were also used. The resistance associated SNPs from iteration 4 (238 SNPs) were also used. | 0.363446 | 0.29105787 |

| | | | | |
|---|---|---|---|---|
| 16 | 14916 | Thinning the genetic dataset by 1 SNP per 15,000 bp window. A minor allele frequency of 0.1 and a mapping quality filter of 15 were also used. The resistance associated SNPs from iteration 4 (238 SNPs) were also used. | 0.4699868 | 0.30724723 |
| 17 (comparison) | 2377 | Using SNPs with a p-value of less than 1E-3 from the GWAS. Genetic data is filtered as described in the manuscript, with a mapping quality filter of 15. | 0.372008 | 0.5069734 |
| 29 | 689 | Thinning iteration 17's SNPs by 1000bp (originally 2377 SNPs) | 0.6344271 | 0.5421888 |
| 33 | 15566 | Using both SNPs from iteration 16 (background SNPs) & 29 (associated SNPs) | 0.6646783 | 0.4503532 |

Source of SNPs

SNPs were predominantly sourced from the GWAS described in the manuscript. To determine the robustness of the GWAS method, an alternative method was also conducted. This method ran a GWAS of the same phenotype and genotype data using a Bayesian Sparse Linear Mixed Model (BSLMM) (Table S4). These results were then used in a genomic prediction model similar to those presented previously. This resulted in similar levels of accuracy.

Epistatic SNPs were also used in our analyses. The inclusion of epistatic SNPs with the strongly associated SNPs, improved model quality. This was unlike a model that used solely epistatically correlated SNPs.

We tested the use of only epistatically correlated SNPs and also the addition of epistatic SNPs in the genomic prediction model. Epistatic SNPs on their own performed poorly as a genomic prediction model. However, the inclusion of the epistatic SNPs in addition to the strongly associated SNPs, improved the model's quality.

*Table S11. The iteration number, the number of SNPs used in the genomic prediction model for the iteration, a description of the SNPs used, the mean reliability of the predictions made and the $R^2$ value of all individuals (training and cross-validation individuals combined)/only cross-validation individuals. Here, the iterations focus on using SNPs from alternative sources, including a GWAS that used BSLMM and epistatically significant SNPs.*

| Iteration | SNP count | Description | Mean reliability | $R^2$ |
|---|---|---|---|---|
| 44 | 1000 | BSLMM top 1000 SNPs | 0.6497708 | 0.3180254 |
| 45 | 500 | BSLMM top 500 SNPs | 0.3650349 | 0.5333694 |
| 40 (comparison) | 996 | Myrtle rust associated SNPs from the original GWAS, thinned at 1 SNP per 100 bp. | 0.6298315 | 0.5697814 |

| | | | | |
|---|---|---|---|---|
| 46 | 997 | Same SNPs as iteration 40, but the bottom 50 SNPs were replaced with SNPs from the BSLMM GWAS that did not overlap. | 0.5446107 | 0.5665020 |
| 43 | 997 | Same SNPs as iteration 40, but the bottom 50 SNPs were replaced with SNPs that were most epistatically associated. | 0.5852423 | 0.5808438 |
